# The Hidden Guardians: From Active Cells to Spores in a 20-Year Journey into the Microbial Heart of a Drying Oasis

**DOI:** 10.1101/2025.10.04.680467

**Authors:** Antonio González-Sánchez, Emmanuel Cordero-Martínez, Dolores Rodríguez-Torres, África Islas-Robles, Norberto García-Miranda, Marisol Navarro-Miranda, Román Zapién-Campos, Felipe García-Oliva, Luis E. Eguiarte, Valeria Souza, Gabriela Olmedo-Álvarez

## Abstract

The Cuatro Ciénegas Basin (CCB) in Coahuila, Mexico, a Ramsar Wetland, is historically characterized by a unique network of pozas renowned for microbial diversity. Among them, the Churince hydrological system epitomized a delicate balance of endemic species, stable water regimes, and complex microbial assemblages. Over the last five decades, however, Churince underwent a catastrophic ∼99% water loss, primarily driven by intensive agricultural extraction. We showed that this desiccation not only caused hydrological collapse but also reshaped microbial physiology. Longitudinal records from 2007 to 2019, combining satellite imagery, piezometer data, and field observations, documented progressive declines beginning in 2011 and culminating in the complete desiccation of the Intermediate Lagoon by 2019. To track microbial responses, we employed a cultivation-based approach distinguishing vegetative cells from dormant spores, enabling direct physiological assessment of metabolic activity versus dormancy. As water diminished, spore abundance increased relative to vegetative forms, revealing a shift toward dormancy as a survival strategy under extreme desiccation. Complementary mesocosm experiments showed that reduced diversity further enriched for spores, illustrating how loss of community complexity accelerated dormancy and erosion of active assemblages. Together, these results demonstrated that habitat loss triggered microbial community change, which we frame as a microbial extension of the ecological concept of community disassembly. By combining two decades of hydrological evidence with microbial physiology, this study fills a critical gap in understanding how wetland drying selects for microbial dormancy. These findings underscore the urgent need for sustainable agriculture and integrated water management to preserve biodiversity and ecosystem function in threatened wetlands worldwide.

## Introduction

Freshwater scarcity is an escalating global threat, reshaping ecosystems across continents. Microbial communities, as the primary drivers of biogeochemical cycles, are essential to ecosystem stability, nutrient recycling, and global resilience. These invisible architects not only sustain critical environmental processes but also represent the evolutionary foundation of biochemical diversity. From large inland lakes to fragile wetlands, the disappearance of water endangers biodiversity, local livelihoods, and ecosystem function.

The Cuatro Ciénegas Basin (CCB), in the Chihuahuan Desert of Coahuila, Mexico, exemplifies this crisis. A globally significant oasis, the CCB is known for extraordinary microbial biodiversity and extreme oligotrophic conditions. Its microbial communities, shaped over geological timescales, provide a unique window into early Earth analogs and microbial adaptation to nutrient limitations. CCB is historically characterized by a unique network of ponds and small lagoons locally called “pozas”, renowned for microbial diversity. Among the aquatic systems in the basin, the Churince hydrological system stood out for its fragile balance and richness in endemic species (Souza *et al*., 2018). Recognizing its significance, it was designated both as a “Protected Area for Flora and Fauna” by Mexican federal government and a Ramsar Wetland of international importance.

The Churince system, situated at a higher elevation than the rest of the basin and bounded by the Sierra de San Marcos y Pinos, historically maintained a stable hydrological regime influenced by ancient groundwater with magmatic signatures. This environment supported gradients in salinity, oxygen, and pH, fostering a mosaic of microhabitats. Previous studies have identified ancient and deeply branching microbial lineages, including the endemic Bacillus coahuilensis, which exemplify adaptations to oligotrophic conditions (Alcaraz *et al*., 2008). Members of the Bacillaceae family, many of which are capable of sporulation, were observed to be abundant in the Churince system (Cerritos *et al*., 2011; Rodríguez-Torres *et al*., 2017), serving as models for microbial survival under nutrient stress.

Over the last five decades, however, the Churince system has undergone severe hydrological collapse, losing more than 99% of its surface water. This desiccation has been driven by unchecked agricultural extraction, especially for the irrigation of alfalfa (Medicago sativa), a highly water-demanding forage crop. In the region, alfalfa is cultivated through two main irrigation practices: flooding of fields using spring water or by center-pivot irrigation systems supplied by wells. Together, these methods withdraw far more water than is replenished annually by precipitation, creating a chronic imbalance between extraction and natural recharge. This imbalance has directly contributed to aquifer depletion and ecological collapse.

Microbial communities are highly responsive to environmental change and can serve as indicators of ecosystem health. Water availability is central to this responsiveness: without extracellular water, nutrients cannot dissolve for cellular uptake, and microbial cells cannot move toward resources, ultimately leading to starvation (Wang & Or, 2013; Schimel, 2018). In drylands, microorganisms often endure prolonged periods of desiccation, facing severe energy limitations and stress. Many responds by entering dormancy, a reversible state of reduced metabolic activity that allows them to survive until favorable conditions return. Dormant microorganisms act as microbial “seed banks,” influencing ecological stability and evolutionary dynamics (Lennon & Jones, 2011). Dormancy has been defined as “a temporary adaptive state of reduced metabolic activity within an extended period of arrested growth that can enable a microbe to maintain viability under unfavorable conditions” (McDonald *et al*., 2024).

Dormancy strategies vary. Some taxa form resting structures with thickened cell walls or extracellular polymers, while others produce spores. Sporulation is well documented in members of Actinomycetota and Bacillota (formerly Firmicutes) phyla, the latter forming endospore, a highly stress-resistant strategy. In this study, we operationally define a shift toward dormancy as a quantifiable increase in the proportion of heat-resistant colony-forming units (CFUs) relative to the total culturable community. Within the Bacillaceae family, exemplified by the model Bacillus subtilis, cells can enter a state of non-replicative viability (quiescence) or initiate sporulation. In the latter case, germination restores vegetative growth when the conditions are favorable (Setlow, 2014; Higgins & Dworkin, 2012). Such Gram-positive phyla are widely distributed in drylands and become ecologically dominant under conditions of desiccation (Leung *et al*., 2020). Accordingly, spore-forming bacteria are also widespread in extreme ecosystems, including deserts (Chanal *et al*., 2006) and endangered salt lakes (Kheiri *et al*., 2023).

Ecological communities are shaped by the dual processes of community assembly, the sequential addition of species over time, and community disassembly, the nonrandom process of progressive species declines and losses (Zavaleta *et al*., 2009, O’Neill, 2016). Community disassembly is not simply the reverse of community assembly, as disassembly is frequently driven by human-caused stressors that were absent when the community originally formed (Zavaleta *et al*., 2009; Gasbarro *et al*., 2019). This process of species loss follows predictable patterns, or “disassembly rules,” where the sequence of decline is governed by species-specific “response traits” that confer vulnerability to stressors such as habitat fragmentation, climate change, or resource depletion. The specific order in which species are lost is critically important, as it can dramatically alter ecosystem functioning, including productivity, nutrient cycling, and the provisioning of ecosystem services (Ostfeld & LoGiudice 2003; Zavaleta *et al*., 2009). While community disassembly is frequently studied in the context of macroorganism responses to human impacts, it also occurs as a natural, cyclical process in ephemeral ecosystems that predictably dry up or decay (O’Neill, 2016; Brown *et al*., 2022).

The sequential disappearance of fish, turtles, and hydrophilic plants in Churince reflects the ecological concept of community disassembly (O’Neill, 2016). Our results extend this framework to microbes: natural habitat loss drive disassembly toward dormancy, and the mesocosms provide a simplified experimental view of the same outcome.

This study arose from a rare, unplanned opportunity to observe microbial responses to real-world environmental collapse. The natural desiccation of the Churince system provided direct insight into how microbial communities shift under sustained drying, unlike experimental models that simulate water loss. By integrating ecological, hydrological, and microbiological data, we address a key gap: no previous study has tracked microbial dormancy through a long-term collapse of a wetland ecosystem.

We conducted a 15-year longitudinal study of the Churince Intermediate Lagoon, using heat sensitivity as a proxy to distinguish between metabolically active (vegetative) and dormant bacterial cells (spores). We quantitatively documented the system’s hydrological decline through water-level measurements, satellite imagery, and field photographs that captured progressive desiccation, including the mortality of fish and turtles. These changes coincided with a marked drop in the water level recorded by the Churince piezometer and were visibly reflected in the surrounding landscape. To complement these observations, we also carried out a two-year mesocosm experiment in which heat treatment reduced community diversity. These simplified mesocosms converged on spore enrichment, reinforcing the idea that lost diversity per se selects for dormancy.

## Materials and Methods

### Study sites, sampling and microbial isolation

The Churince hydrological system, located at 730 m above sea level, comprises a Spring (S), an Intermediate Lagoon (IL), and a Desiccation Lagoon (DL), interconnected by a small river. In the IL, longitudinal sampling was conducted at eight shoreline sites (Li1–Li8, 30–50 m apart) in 2011, 2012, 2019, 2023, and 2025. In 2012, a transect across the IL included sediments collected every 50 m along 500 m eastward (grassland) and 550 m westward (*sotol* dominated (*Dasylirion))* from the shore, yielding 40 samples. In 2014, a 60 cm sediment core was taken at Li7 and sectioned into 2 cm intervals, with 10 subsamples analyzed. GPS coordinates for all sites are provided in Supplementary Tables 1–2.

For longitudinal and transect samples, surface sediments (∼1 cm) were collected with sterile 50 mL tubes. Approximately 0.1 g of sediment was serially diluted (up to 10⁻⁵) in PBS and plated on Marine Medium agar. Core subsamples (0.5 g mL⁻¹) were resuspended in Tris-HCl buffer (pH 8.4), agitated (600 rpm for 1 h), diluted, and plated similarly.

In 2014, when surface water was still present, sediments from three IL sites (Li4, Li5, Li8) were also used to establish laboratory mesocosms. Each sample was divided into two treatments: untreated (intact communities) and heat-treated (selecting for heat-resistant spores). Triplicate mesocosms were maintained in glass flasks at 20 °C under a light–dark cycle, with water added only to maintain sediment coverage. Subsamples were taken periodically over two years, and total CFU and heat-resistant fractions were quantified by plating on Marine Medium.

All samples were collected with federal permission (permit SGPA/DGVS/04512) and transported (at 4°C) either to the CBTA22 laboratory in Cuatro Ciénegas or to the CINVESTAV Irapuato laboratory for plating.

### Colony count and Sporulation percentage

In most sampling schemes, CFU counts were obtained under two conditions: (1) Direct plating, which recovers colonies arising from both vegetative cells and spores that germinate upon plating; and (2) Heat-treated samples, in which sediments were heated at 80 °C for 30 min to inactivate vegetative cells and selectively recover thermotolerant bacteria or spores that readily germinate and grow into colonies. The plating workflow and cell cycle are illustrated in Fig. 1. Sporulation percentage was determined by: (CFU from spores / Total CFU) × 100. Plates were incubated at 28 °C for 5 days.

**Fig. 1.**
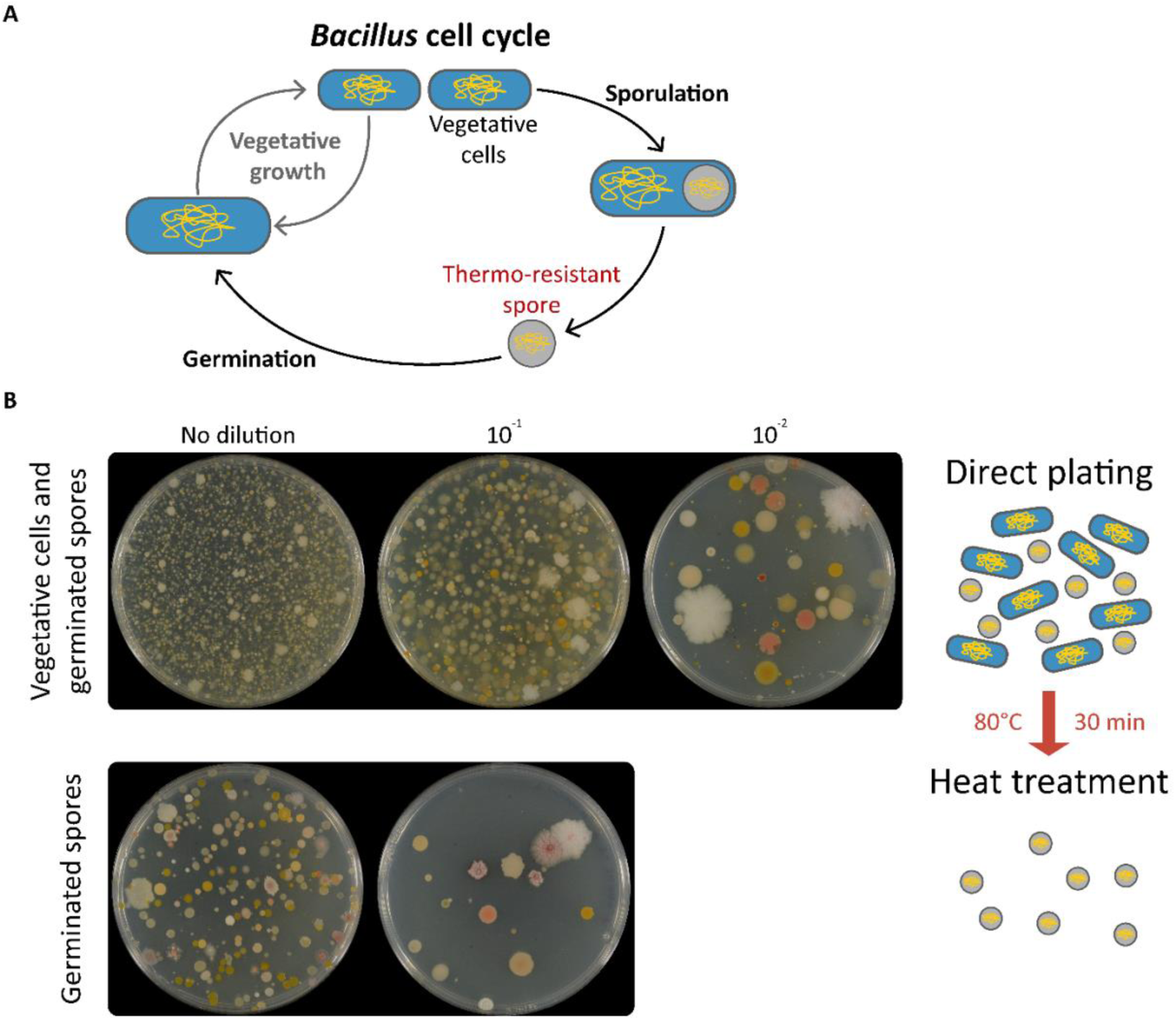
Detection of vegetative cells and spores by direct and heat-treated plating. Representative colony counts obtained from two cultivation approaches. Direct plating recovers both vegetative cells and spores, thereby reflecting the total viable count. Heat-treated plating (80 °C, 30 min) inactivates vegetative cells, allowing only heat-resistant spores to germinate. In this example, direct plating at the 10⁻³ dilution yielded ∼3.1 × 10⁵ CFU, whereas heat-treated plating at the 10⁻¹ dilution yielded ∼2 × 10³ CFU, corresponding to <1% sporulation. Isolates were plated on Marine Medium and incubated at 28 °C for 5 days. The example corresponds to a 2019 sediment sample. A schematic of the Bacillus life cycle is included, showing binary fission under favorable conditions and sporulation under stress.

Cell counts from the 2007 sampling were not obtained because the bacterial isolation methods used that year focused on recovering heat-resistant bacteria (See Supplementary Table 1). For the vertical depth profile study, colonies were additionally classified into 19 morphotypes based on colony shape, margin, elevation, texture, and color (Supplementary Table 3).

### Satellite imagery and water level measurements in the Churince hydrological system

Satellite imagery was obtained from Google Earth (7.3.6.10201) using the time-lapse feature to access historical images. Water levels in the Churince Spring were monitored from 2007 to 2015 using a piezometer installed in a borehole by the Comisión Nacional de Áreas Naturales Protegidas (CONANP). Water level was recorded as the vertical distance between the piezometric sensor at the borehole bottom and the water surface above the sensor. Measurements were taken in different months each year (Supplementary Table 5). In 2012, a temporary disconnection in monitoring occurred. The lowest recorded water level was observed in 2015, marking the end of the measurement period.

### DNA extraction

A total of 1,419 bacterial isolates were obtained from samples of 2007, 2011, 2012, 2014, 2019, and 2022. Total DNA was extracted for PCR amplification of the 16S rRNA gene using a modified phenol–chloroform method with bead beating (0.1 mm glass beads) to lyse cells. Briefly, overnight cultures were centrifuged (16,000 × g, 5 min), washed with TE buffer, and resuspended in lysis solution (2% Triton X-100, 1% SDS, 100 mM NaCl, 10 mM Tris–HCl pH 8.0, 1 mM EDTA). After the addition of TE buffer and phenol–chloroform, 0.1 mm glass beads were incorporated, and samples were vortexed for 5 min to enhance cell disruption. The aqueous phase obtained after centrifugation was re-extracted with chloroform, and DNA was recovered by ethanol precipitation, washed with 75% ethanol, air-dried, and resuspended in nuclease-free water containing RNase (2 μg/mL).

### Taxonomic classification and phylogenetic analysis

Taxonomic classification was based on 16S rDNA sequences amplified using universal primers 27F (5′-AGAGTTTGATCCTGGCTCAG-3′) and 1492R (5′-TTACGGYTACCTTGTTACGACT-3′). PCR products were sequenced by Sanger technology at LANGEBIO-CINVESTAV, Irapuato, Mexico. A total of 1,419 full-length 16S rDNA sequences were obtained and deposited in the NCBI database; accession numbers are listed in Supplementary Table S6. Taxonomic assignment of each isolate was performed using BLASTn against the NCBI 16S ribosomal RNA database, complemented with SILVA classification for confirmation.

For phylogenetic analysis, the 1,419 sequences were trimmed to ∼350 bp and aligned with the ClustalW algorithm. Phylogenetic reconstruction was conducted in RAxML-GUI using the Maximum Likelihood (ML) method with the K2+G evolutionary model and 1,000 bootstrap replicates. Fusobacterium was used as the outgroup. The resulting tree was visualized and annotated in the Interactive Tree Of Life (iTOL) platform (Letunic and Bork, 2024).

### Statistical Analyses of CFU, Sporulation, and Community Structure

Differences in CFU counts and sporulation percentages among the longitudinal time series, spatial transect, and vertical depth profile datasets were evaluated using one-way analysis of variance (ANOVA), with a significance threshold of α = 0.05.

Microbial community analyses were conducted for the classified 1,419 bacterial isolates. For longitudinal comparisons, isolates were grouped into three ecological phases representing distinct hydrological states of the Churince System: With Water (2007), Transition (2011, 2012, 2014), and Desiccation (2019, 2022) (Supplementary Table 6). Community composition differences were assessed via Permutational Multivariate Analysis of Variance (PERMANOVA, α = 0.05) applied to both alpha and beta diversity indices. Dissimilarity matrices were computed using Bray–Curtis (abundance-based) and Jaccard (presence/absence-based) distances. Beta diversity patterns were visualized using Principal Coordinates Analysis (PCoA). All community-level statistical analyses were performed in R using the vegan package, with visualizations generated in ggplot2 (Supplementary Figs. 4 and 5).

For genus-level analyses, taxonomic assignments were aggregated at the genus level, and both relative and absolute abundances were calculated as the number of isolates per sampling site within each ecological phase (Supplementary Table 7). Isolates belonging to Bacillus were examined separately (Supplementary Table 8 and Supplementary Fig. 6). Differences among phases were tested with the Kruskal–Wallis test (α = 0.05), statistic (H) and its associated p-value indicated whether at least one group differed significantly in distribution (Supplementary Table 9). This analysis was performed in Python 3.11 using the kruskal function from scipy.stats module.

## Results

### Hydrological Collapse of the Churince Intermediate Lagoon and Agricultural Drivers

The Cuatro Ciénegas Basin (CCB) exemplifies the fragility of desert wetlands. Within it, the Churince hydrological system—comprising the Desiccation Lagoon (DL), the Intermediate Lagoon (IL), and a Spring (WS)—was once an oasis of exceptional biodiversity (Souza *et al*., 2018), home to endemic fauna such as *Eleutherodactylus* sp. nov., *Terrapene coahuila*, and *Cyprinella xanthicara* (Supplementary Fig. 2). Photographs from 2001 and satellite imagery from 2009 confirm that the Intermediate Lagoon still retained water at the beginning of our study (Fig. 2).

**Fig. 2.**
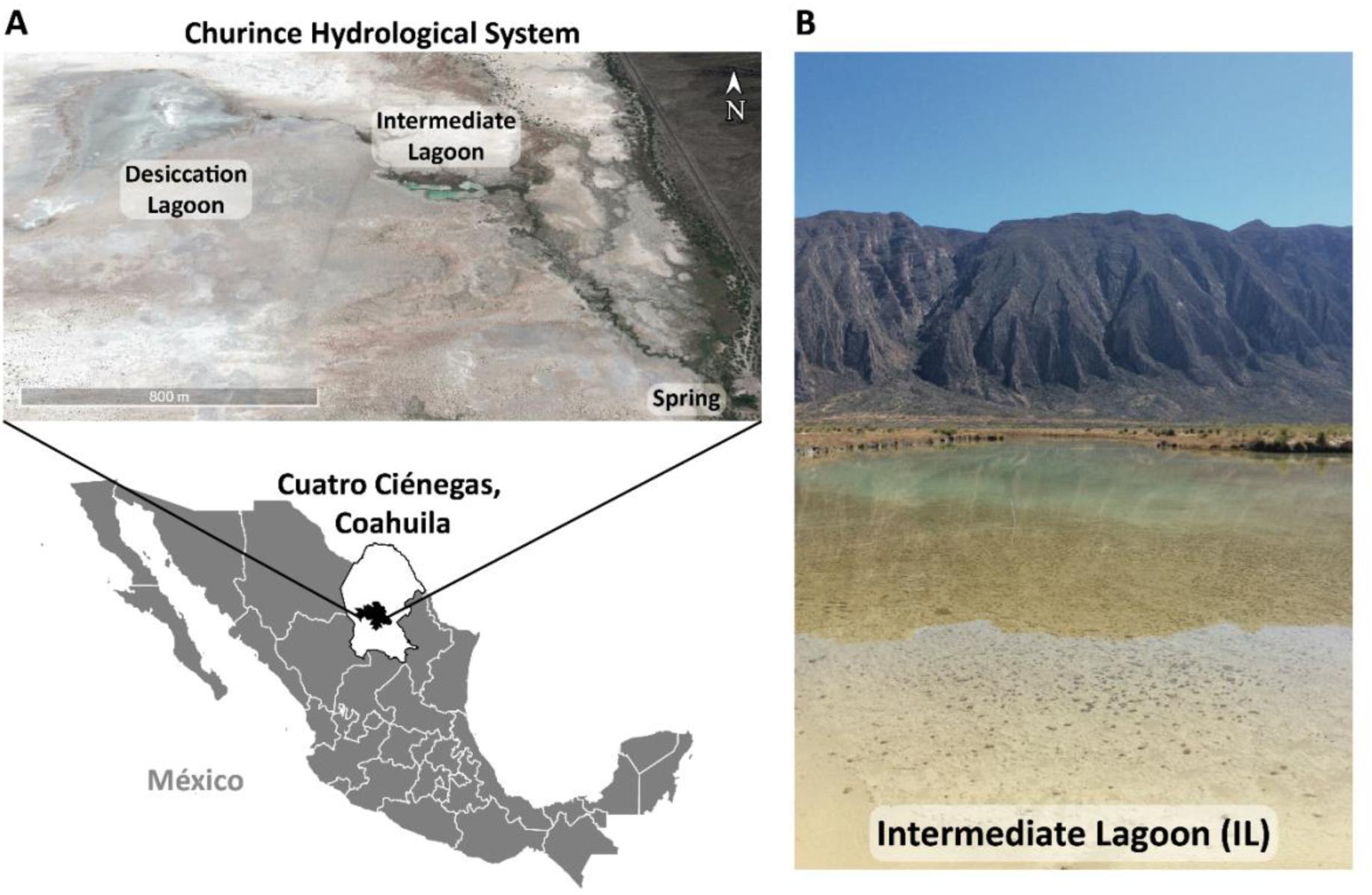
The Churince System in Cuatro Ciénegas, Coahuila, México. A) The Churince system consists of the Desiccation Lagoon (DL), Intermediate Lagoon (IL), and Spring (S), connected by an arroyo. The satellite image was taken from Google Earth from 2009, when the lagoons still held water. B) Photograph of the Intermediate Lagoon in 2001, when the system still retained surface water.

Multiple lines of evidence converged on a rapid collapse. Satellite images revealed the water loss: abundant water in 2009, drastically reduced by 2012, and gone by 2019 (Fig. 3A). The piezometer records (2007–2015) showed a steady groundwater decline (Fig. 3B), with a sharp drop in 2011 coinciding with observations of dead turtles and fish carpeting exposed sediment (Supplementary Fig. 3). Photographs from 2014 and 2019 further documented receding shorelines and wildlife loss. By 2019, the Intermediate Lagoon had dried completely.

**Fig. 3.**
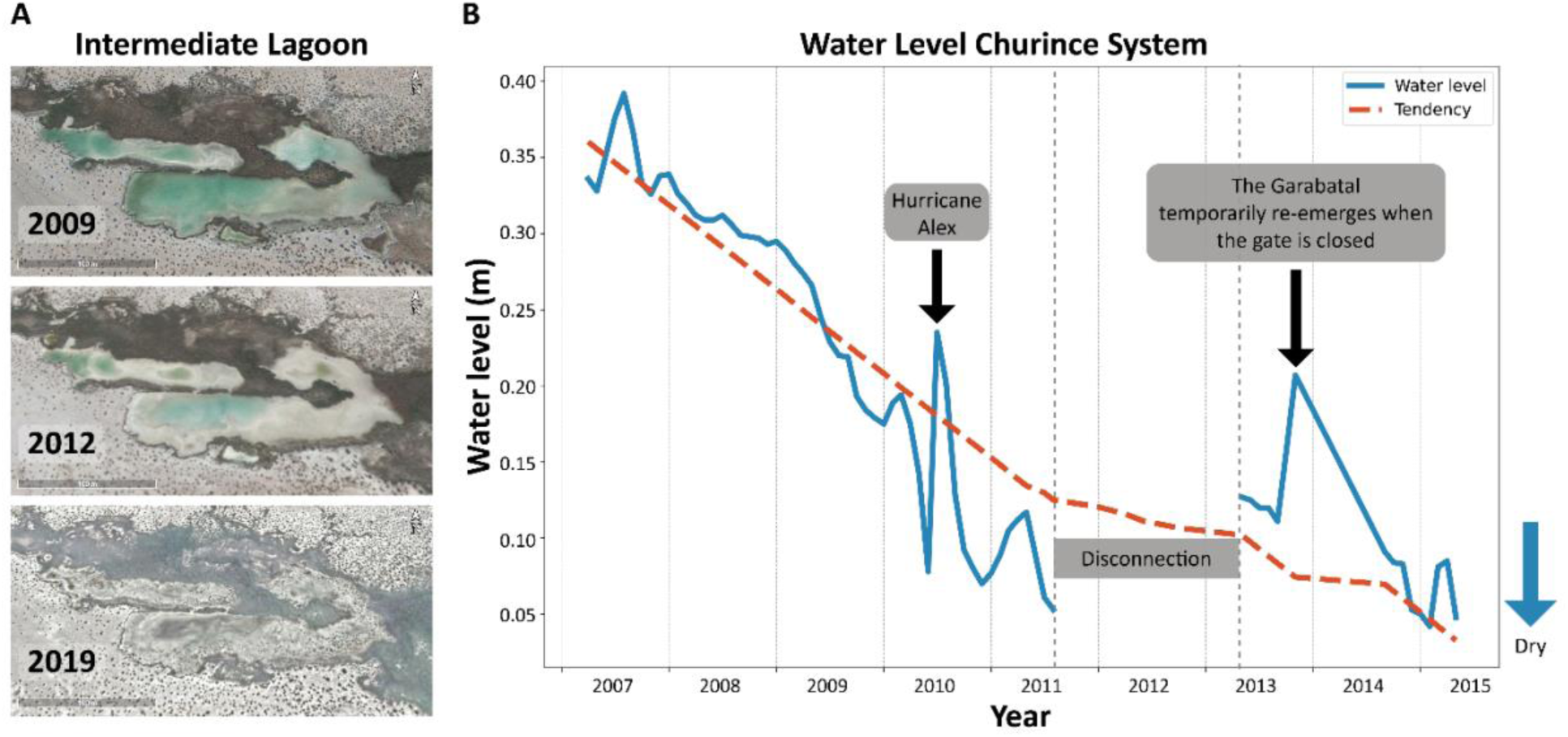
Gradual desiccation of the Intermediate Lagoon in the Churince system. (A) Satellite images (Google Earth) showing progressive water loss and increasing aridity in the Intermediate Lagoon and surrounding area: 2009 (with water), 2012 (reduced water), and 2019 (completely dry). (B) Yearly average groundwater levels (2007–2015) measured by a piezometer installed at the Churince well. The piezometer records the height of the groundwater table relative to the surface, reflecting the water available to feed the lagoon. A transient rise in 2010 corresponds to Hurricane Alex and the brief re-emergence of the Garabatal spring after the temporary closure of the gate. Piezometer data are provided in Supplementary Table 5.

This collapse coincided with intensified agriculture (Supplementary Fig. 1). Regional water extraction rose from 22 million m³ in 1997 to 88 million m³ in 2022 (public statistics: IMTA, 2023; compiled from CONANP monitoring), driven largely by alfalfa (Medicago sativa), a water-demanding forage crop. In the nearby Hundido Valley, alfalfa is cultivated through flood irrigation and center pivots, consuming ∼15,660 m³/ha. Satellite images show both the reduction of surface water in Churince and expansion of alfalfa fields, from 21 active plots in 2007 to 29 in 2020 (Supplementary Fig. 1B). Despite its Ramsar designation, agricultural expansion strongly overlapped with and likely drove the hydrological collapse of the Intermediate Lagoon.

### Shifts in dormant vs. active bacterial fractions

Following the disassembly of the macrofaunal community (Supplementary Fig. 3), we analyzed microbial responses to desiccation. Sediments cultured on Marine Medium consistently recovered Bacillaceae. Heat-treated samples revealed the spore-forming fraction, while untreated samples reflected the total culturable pool (vegetative cells plus germinable spores, detailed in Fig. 1). The proportion of heat-resistant CFUs served as a proxy for dormancy. To capture ecological patterns, we used three strategies: a vegetation transect, a vertical sediment core, and a shoreline series spanning more than a decade of decline.

In the transect (2012, Fig. 4C), sporulation co-varied with vegetation and aridity: on the sotol-dominated west side nearly all colonies (∼100%) originated from spores, whereas the grass-covered east side showed ∼25% (Fig. 4B). Total CFU counts were ten times higher in grassland soils (∼25,000 CFU/mL), indicating a larger active fraction in grassland, while spores dominated in arid zones (Fig. 4; Supplementary Table 2).

**Fig. 4.**
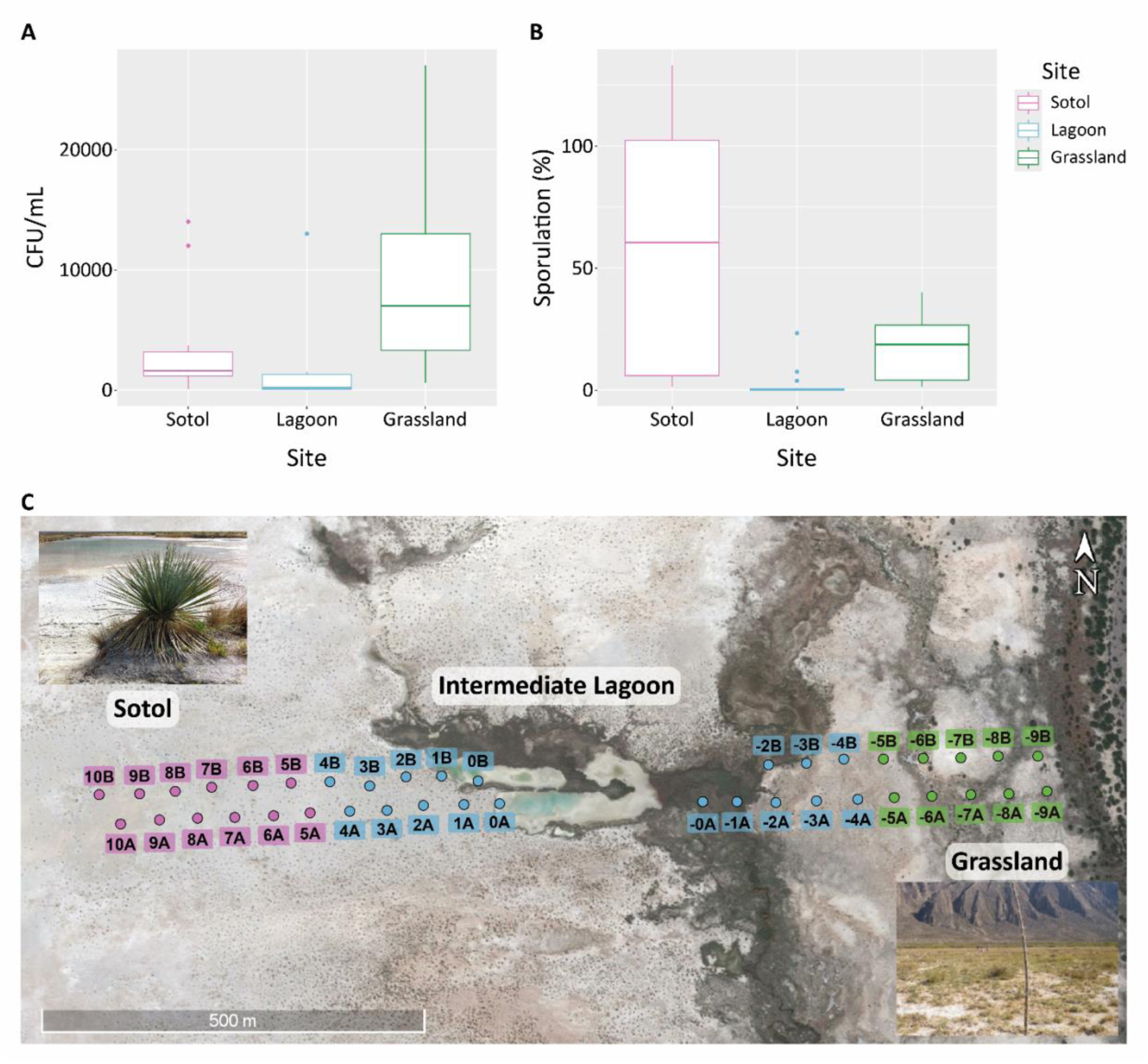
Transect sampling of the Churince System (2011). (A) Average colony-forming unit (CFU) counts per site, with the highest values recorded in grassland samples. (B) Sporulation percentages along the east–west transect highest at sotol sites, lowest at lagoon sites, and intermediate at grassland sites. (C) Google Earth image (2012) showing transect sampling points at 50 m intervals: pink squares denote sotol (*Dasylirion*) sites, blue squares lagoon sites, and green squares grassland sites. Each site had a replicate 20 m away, labeled A and B. Representative photographs of sotol (left) and grassland (right) landscapes are shown for context.

The 2014 sediment core (Fig. 5A) represents decades of deposition. Despite burial, vegetative cells were consistently detected across all layers, where the total CFU always exceeded spore CFU (Fig. 5B). Higher counts were generally observed in the deeper, moister sections (Fig. 5B; Supplementary Table 4). Although the core appeared homogeneous, taxonomic composition and colony morphotypes differed among layers, indicating persistent microbial activity and structuring at depth (Fig. 4C). The relative abundance of the sequenced isolates (n=42) shows that nearly 40% belong to the Bacillus genus (Fig. 4D, Supplementary Table 6).

**Fig. 5.**
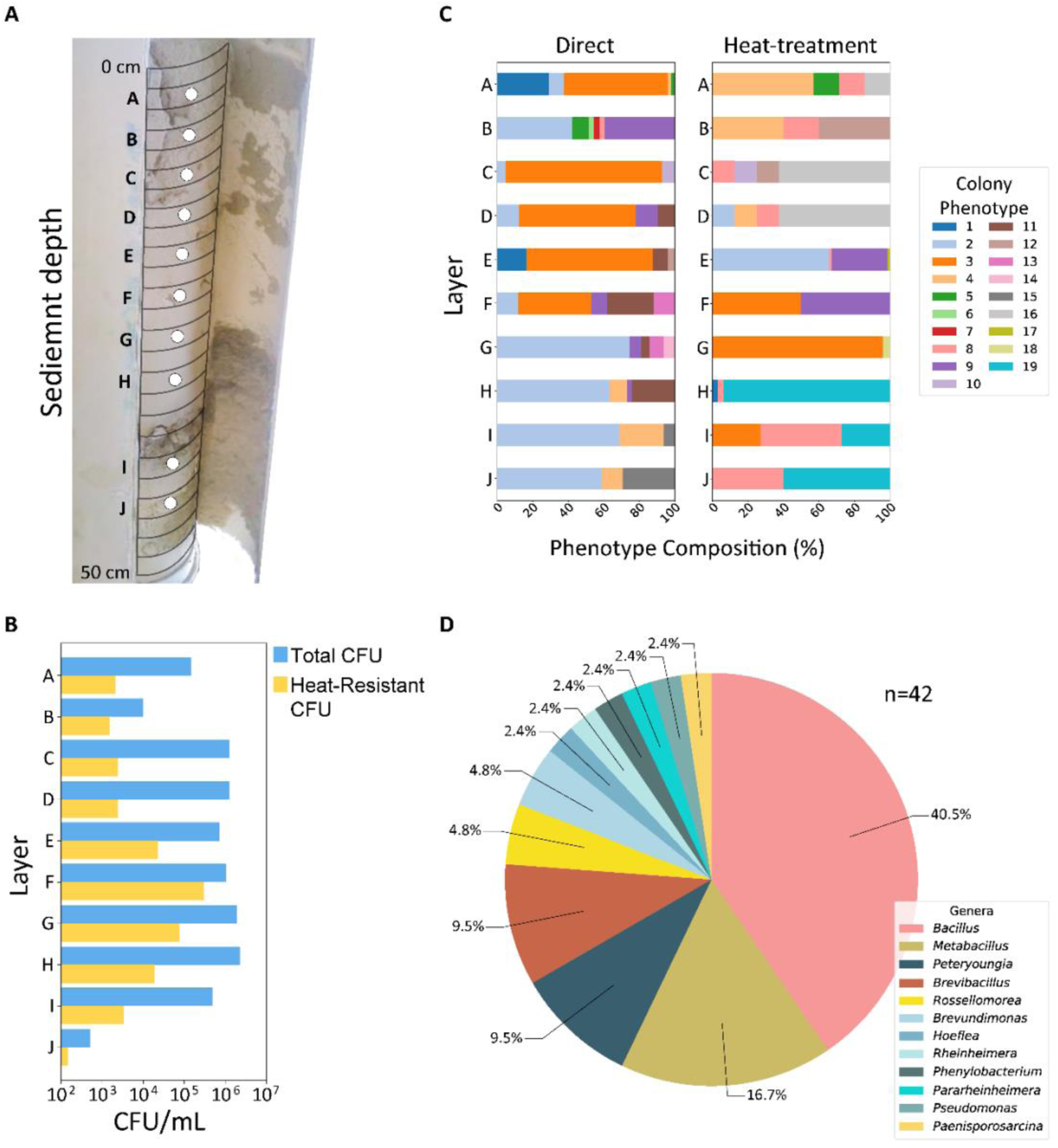
Vertical core sampling at point IL7 in the Intermediate Lagoon. (A) Photograph of a PVC core (cut lengthwise) collected at IL7, sectioned into 2 cm segments, with ten sampling points (A–J) marked by white circles. (B) Phenotypic diversity of isolates from each layer: left panels show direct plating (without heat treatment), right panels show phenotypes after heat treatment. (C) Colony-forming unit (CFU) counts per layer: blue bars, direct plating; yellow bars, heat-treated counts. (D) Taxonomic diversity of 41 isolates based on 16S rDNA sequencing. The pie chart shows genus-level relative abundances, with Bacillus as the predominant genus.

The shoreline series (2011–2025, Fig. 6A) documented a progressive shift toward dormancy. In 2011 and 2012, fewer than 5% of colonies originated from spores. By 2019, sporulation increased to 10–20%. In 2023, it exceeded 20% at some sites, and by 2025, it reached an average above 30% (Fig. 6C; Supplementary Table 1). This trajectory represents a trait-mediated community disassembly, where Bacillaceae shifted from vegetative states into dormancy as surface water disappeared.

**Fig. 6.**
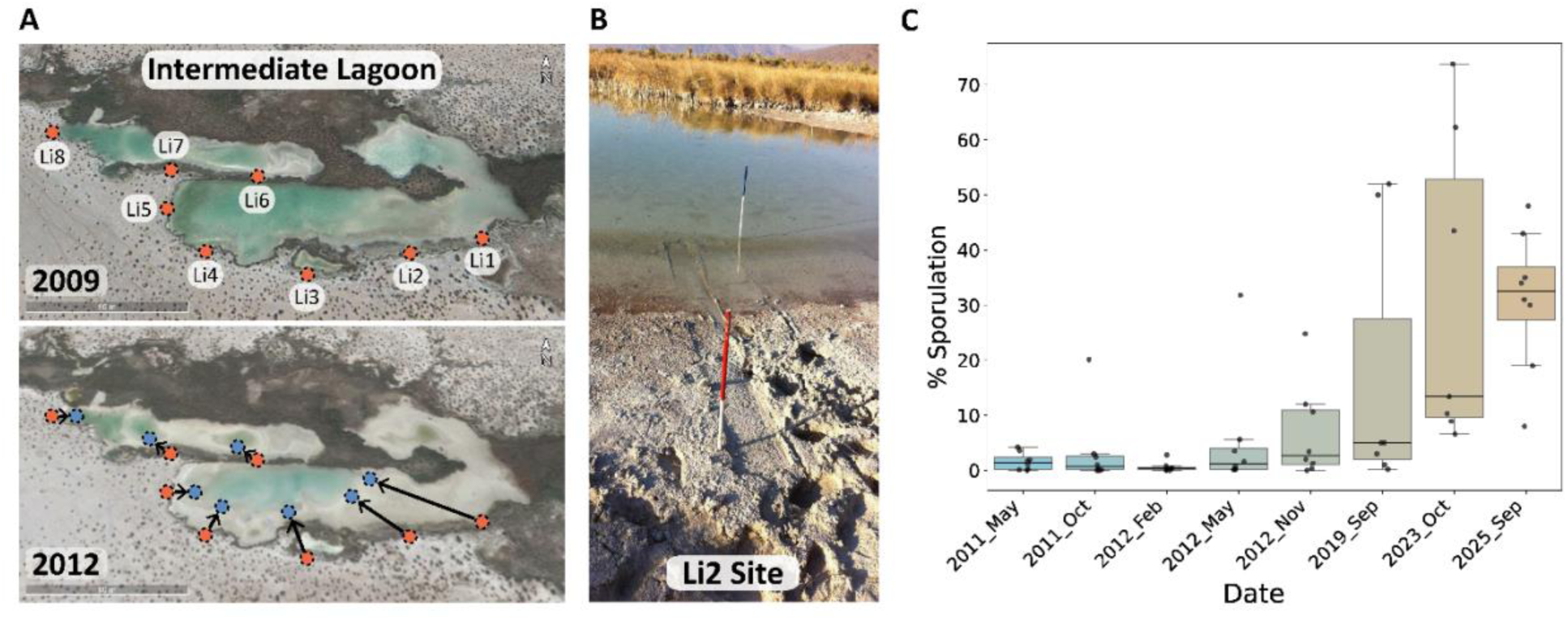
Longitudinal analysis of microbial activity during water loss in the Churince Intermediate Lagoon (2011–2025). (A) Google Earth images from 2009 and 2012 showing sampling points (Li1–Li8) established to collect sediments annually along the shoreline while it was submerged. As the water receded, new sampling poles were installed progressively toward the center of the lagoon to continue sampling submerged sediments. (B) Photograph of the Intermediate Lagoon 2011 showing a red sampling tube located in now-dry sand near the original shoreline and a blue tube placed farther into the lagoon, indicating pole relocation as water levels declined. (C) Sporulation percentages in sediment samples over time (2011, 2012, 2019, 2023 and 2025), showing a steady increase as the lagoon dried. Values represent the average across all sampling points for each date.

### Composition of culturable bacteria

The drying of Churince was reflected not only in activity fractions but also in community composition. Across all sampling years, we sequenced and classified 1,419 bacterial isolates from the Intermediate Lagoon (Fig. 7A), of which 682 belonged to the genus Bacillus (Supplementary Tables 6 and 7). At the genus level, isolates were grouped into three ecological phases: With Water, Transition, and Desiccation (Fig. 7B), corresponding to different years (see Methods).

**Fig. 7.**
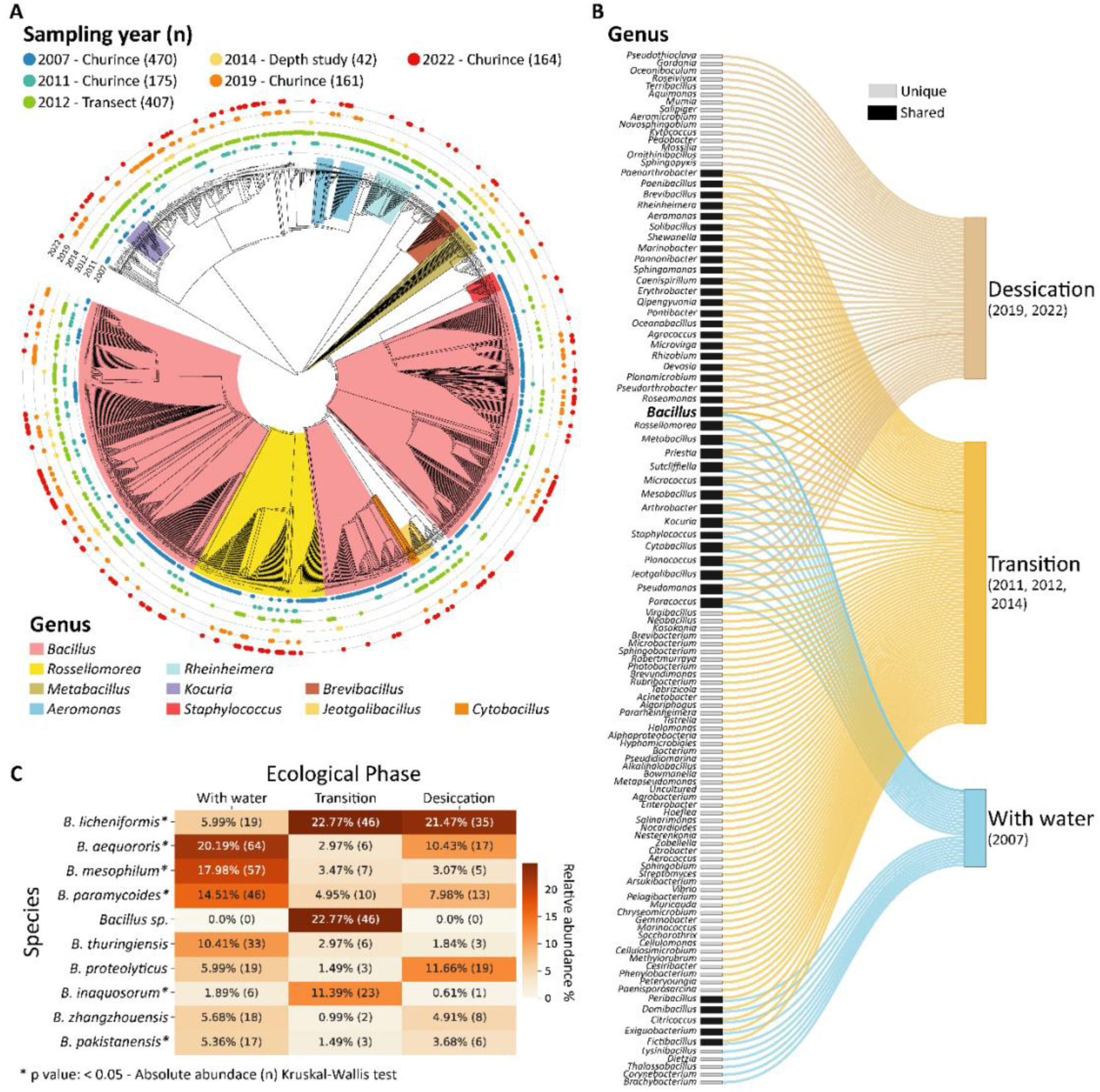
Phylogenetic diversity, taxonomic composition, and ecological phase patterns of bacterial isolates from the Churince System. (A) Maximum Likelihood phylogenetic tree based on partial 16S rRNA gene sequences (∼350 bp) from 1,419 isolates, color-coded by taxonomic affiliation and sampling year. Numbers of isolates per year are shown in parentheses. (B) Sankey diagram showing the distribution of genera across three ecological phases corresponding to distinct hydrological states: With Water, Transition, and Desiccation. Grey boxes denote genera unique to a single phase; black boxes denote genera present in at least two phases. (C) Heatmap of the ten most abundant *Bacillus* species across the three phases, with absolute abundances in parentheses. Statistical differences among phases were tested using the Kruskal–Wallis test. The complete distribution of *Bacillus* species and statistical analysis is provided in Supplementary Tables 8 and 9.

From these ecological phases, beta diversity was assessed using PCoA based on Bray– Curtis and Jaccard distances, which showed marginal, non-significant separation among phases (Bray–Curtis: R² = 0.47, p = 0.267; Jaccard: R² = 0.42, p = 0.267) (Supplementary Fig. 4). Nevertheless, genus-level composition shifted across phases (Fig. 7B).

Alpha diversity revealed an unexpected pattern: the water phase had the lowest diversity, likely due to the 2007 focus on heat-resistant bacteria, which underestimated the number of vegetative cells. The transition phase exhibited the highest Shannon and Simpson values, indicating greater richness and evenness of the species composition. The Desiccation phase maintained high richness, probably due to rare dormant lineages, but with reduced evenness (Supplementary Fig. 5). These results suggest that transitional conditions supported more complex communities than either extreme.

Despite these shifts, a core group of taxa persisted across all phases, including several spore-forming Bacillaceae (*Bacillus, Rossellomorea, Metabacillus, Priestia, Sutcliffiella, Mesobacillus, Cytobacillus,* and *Jeotgalibacillus*) as well as *Micrococcus, Arthrobacter, Kocuria, Pseudomonas, Planococcus,* and *Paracoccus* (Fig. 7B). Within this core, *Bacillus* represented 67.45%, 32.37%, and 50.15% of isolates in the water, transition, and desiccation phases, respectively (Supplementary Table 7). The elevated proportion observed in the water phase likely reflects a methodological bias, as isolations at that time relied on heat-treated samples, favoring spore-forming and thermotolerant taxa. Nonetheless, *Bacillus* abundance increased again from the transition to desiccation phases, reaching ∼50% of isolates.

Species-level analysis showed that *B. licheniformis* rose from ∼6% in 2007 to ∼21% in 2022, while species such as *B. aequororis, B. paramycoides*, and *B. mesophilum* declined with drying (Fig. 7C; Supplementary Tables 8–9). Thus, surface water loss shifted the culturable community toward spore-forming *Bacillus* species tolerant of desiccation (Supplementary Fig. 6).

### Reduced community diversity drives microbial assemblages toward dormancy

Field data showed habitat loss, diversity loss, and desiccation occurring simultaneously. To isolate the specific effect of community simplification, we established a two-year mesocosm experiment using lagoon sediments, which were maintained under a light–dark cycle and watered only to offset evaporation. Treatments compared intact communities with those exposed to an initial heat treatment that eliminated non-spore-forming heterotrophs and primary producers (cyanobacteria and algae).

Our results showed a sharp divergence. Untreated mesocosms, which retained a complete microbial food web, maintained predominantly active communities, with spores representing only 8–27% of culturable bacteria (Fig. 8A; Supplementary Fig. 7A). In contrast, heat-treated mesocosms, reduced to heterotrophs, rapidly shifted toward dormancy: spores accounted for 36–46% of CFU (Fig. 8B; Supplementary Fig. 7B). Heat treatment caused substantial mortality, releasing necromass that fueled a short-lived bloom of endospore-forming Bacillaceae, followed by collapse into spore dominance. Colony morphotypes confirmed that spore-former diversity was retained even as autotrophs and competitors were lost. Vegetative and spore counts showed mirror trajectories, with about half of the surviving bacteria sporulating soon after the perturbation (Supplementary Table 10).

**Fig. 8.**
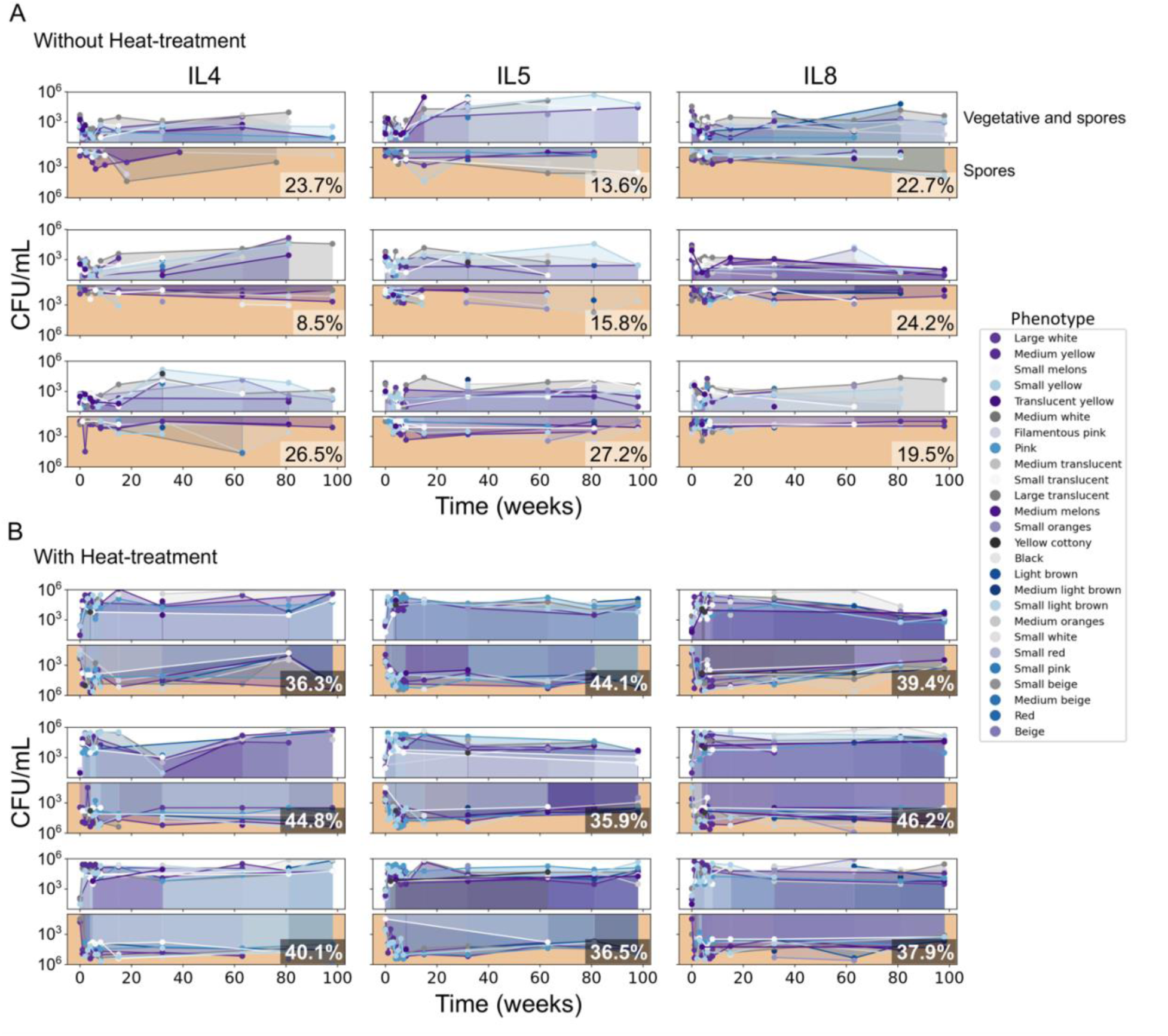
Population dynamics of bacterial phenotypes under direct and heat-treated mesocosm conditions. Mirror plots show temporal changes in colony-forming units per milliliter (CFU/mL). In each plot, the **upper segment** shows total counts from direct plating (vegetative cells + spores), and the **lower segment** (orange background) shows counts after heat treatment (80 °C, 30 min; spores only). Each line with shaded area denotes a distinct bacterial phenotype, consistently color-coded across treatments. Columns correspond to site samples IL4, IL5, and IL8 (see Fig. 6), with three independent replicates per site (nine plots per panel). Mean sporulation (%) across all sampling times shown in the bottom right of each plot. Panel (A) shows mesocosms initiated without prior heat treatment; panel (B) shows mesocosms initiated with heat treatment..

Together, these results suggest that community disassembly, characterized by the non-random loss of autotrophs and heat-sensitive heterotrophs, drives ecosystem collapse. The experimental trajectory mirrors natural desiccation in Churince, where removal of functional groups eliminated nutrient inputs and forced the community into dormancy.

## Discussion

The desiccation of the Churince system illustrates the ecological consequences of chronic groundwater overexploitation. Surface water loss paralleled groundwater decline (Fig. 3), consistent with global patterns of aquifer depletion in arid regions under irrigation pressure (Bierkens & Wada, 2019). In the Cuatro Ciénegas Basin, alfalfa cultivation has been a major driver of extraction (Ortiz Acosta & Romo Aguilar, 2016; Souza *et al*., 2006). Despite its Ramsar designation, the Intermediate Lagoon disappeared by 2019, underscoring governance failures and the limits of international protection (Davidson *et al*., 2023). Satellite imagery and field photography corroborated this trajectory, documenting shoreline retreat and mass mortality of fish and turtles. The sequence of losses, from macrofauna to microbial shifts favoring spore-formers, reflects a non-random process shaped by desiccation. Ecologists describe this unraveling as community disassembly (Zavaleta *et al*., 2009; O’Neill, 2016). While the system is not naturally ephemeral, anthropogenic stress forced a rapid decline resembling ephemeral habitats. Pisanty *et al*. (2022) similarly documented riparian plants in Churince as “canaries,” foreshadowing collapse. Here, the same logic applies hierarchically, from vertebrate loss to microbial shifts dominated by spore-forming Bacillaceae.

Our assays highlight the central role of Bacillaceae, allowing spores to serve as proxies for declining metabolic activity. Across a vegetation transect, shoreline series, and sediment core, the narrative was consistent: aridity reduces microbial activity and favors dormancy, while residual moisture sustains active cells. In the core, subsurface layers retained water and supported higher proportions of vegetative cells, paralleling reports of viable and metabolically active cells persisting for decades in lake sediments (Haglund *et al*., 2003). These findings indicate that spore banks in Cuatro Ciénegas are not inert repositories but dynamic reservoirs, where water and porosity may allow oxygen and nutrients to sustain metabolic activity even after long-term burial.

Taxonomic shifts reinforce this interpretation. A core group of spore-forming Bacillaceae persisted, and within the genus *Bacillus,* some species such as *B. licheniformis* increased, while others, including *B. aequororis, B. mesophilum* and *B. paramycoides*, declined. This pattern highlights how species composition changes across ecological phases. Alpha diversity patterns illustrate the trajectory: stable aquatic conditions supported few dominant taxa, transitional phases maximized richness and evenness, and desiccation maintained high richness—likely due to dormant lineages—although evenness decreased. Similar enrichment of spore-formers has been reported in deserts (Chanal *et al*., 2006; Piubeli *et al*., 2015; Ramos-Madrigal et al., 2024) and endangered salt lakes (Abdallah *et al*., 2016; Kheiri *et al*., 2023; Stulina *et al*., 2019), underscoring global relevance. Spores are known to survive on geological timescales (Vreeland *et al*., 2000), and in Cuatro Ciénegas they may preserve lineages otherwise lost to desiccation.

Mesocosm experiment adds mechanistic insight: disassembly altered energy flow. Untreated systems, with autotroph–heterotroph linkages, sustained nutrient cycling and photosynthesis. Heat-treated systems lost this linkage, becoming dependent on necromass from the die-off. Thus disassembly, whether driven experimentally or environmentally, pushes communities toward persistence via dormant spores rather than active cycling.

Dormancy provides both refuge and constraint: it preserves rare taxa but reduces contributions to nutrient cycling (Allison & Martiny, 2008; Prosser *et al*., 2007; Lennon & Jones, 2011). Sporulation may serve as a marker of resilience and an early warning of decline. Our mesocosm results refine the “microbial seed bank” hypothesis (Lennon & Jones, 2011; Shoemaker & Lennon, 2018). While spores remain viable, they cannot by themselves re-establish complex nutrient flows. Self-sustaining function requires phototrophs, heterotrophs, and higher-order interactions. Without diverse guilds, spores remain dormant even when water returns. Thus seed banks preserve diversity, but cannot guarantee recovery. Future recolonization will likely involve new organisms, reshaping the community.

Our cultivation-based approach recovered only a subset of microbial diversity, specificaly organisms capable of growing aerobically on Marine Medium. Although not selective for Bacillota, this medium has been shown to support their growth and efficiently recover diverse taxa, including Bacillota and Actinomycetota (Rodrigues & Carvalho, 2022), thus enabling detection of both active cells and dormant forms. Nevertheless, many inactive bacteria likely rely on alternative survival strategies that were not captured in this study. Molecular methods provide complementary views: DNA vs. RNA comparisons identify active fractions, and functional markers such as *spo0A* track endospore formers (Filippidou *et al*., 2016). Future research integrating omics and physiology of Bacillaceae will identify genes critical for survival under desiccation. Continued monitoring will be essential, especially if restoration efforts occur, to evaluate the potential for seed banks to reestablish function. Exploring alternative strategies, such as the VBNC state, will also be key (Setlow & Christie, 2023).

At a broader scale, the disappearance of Churince provides a rare longitudinal view of microbial responses to wetland drying. It shows that desiccation extinguishes macrofauna and reconfigures microbes toward dormancy. This signal highlights the urgent need for integrated water governance where agriculture and climate variability intersect. Simple assays quantifying active versus dormant bacteria can reveal tipping points when resilience erodes and irreversible transitions become imminent. These microbial signals parallel the theory of critical slowing down (Scheffer *et al*., 2001; Scheffer *et al*., 2012). In Churince, they were recognized only after collapse. This underscores the importance of monitoring microbial assays in conjunction with hydrology and chemistry. The collapse also highlights the importance of incorporating microbes into conservation policy. The launch of the IUCN Microbial Conservation Specialist Group (Gilbert *et al*., 2025) marks a milestone, and our results provide a case of why it matters: microbial communities signal decline and embody resilience that must be protected.

## Conclusion

Desiccation of the Churince Intermediate Lagoon exemplifies hierarchical community disassembly, from vertebrate loss to trait-mediated microbial collapse. Driven by unsustainable groundwater use, it also highlights a practical tool: simple microbial assays as low-cost early-warning indicators of wetland collapse. Longitudinal data indicates that aridity induces endospore-forming Bacillota into dormancy and reduces surface activity. Spores function both as long-term survival structures and, for the first time, as a measurable proxy for metabolic activity under desiccation. This response combines resilience and loss: seed banks preserve rare taxa yet signal reduced activity and ecosystem function. Because microbes underpin nutrient cycling, energy flow, and trophic stability, their collapse represents a fundamental functional loss, not merely a taxonomic shift. Overall, community disassembly spans macro to microorganisms, and microbial responses, both revealing and driving functional decline, strengthen the case for integrated water governance to safeguard desert wetlands.

## Acknowledgments

We thank SEMARNAT for granting permission to sample in the Cuatro Ciénegas Basin Natural Protected Area (permits SGPA/DGVS/04512; SBRA/DGVS/02870/25); CONANP for providing access to piezometer data; and Churince UMA for granting site access. We thank Manuel García-Ulloa and Jaime Gasca Pineda for their valuable help during this work. We also thank the many undergraduate, graduate, and postdoctoral students who contributed to the sampling and technical laboratory work. Graphic abstract by Enrique Hurtado-Bautista.

## Supplementary figures

**Supplementary Fig. 1.**
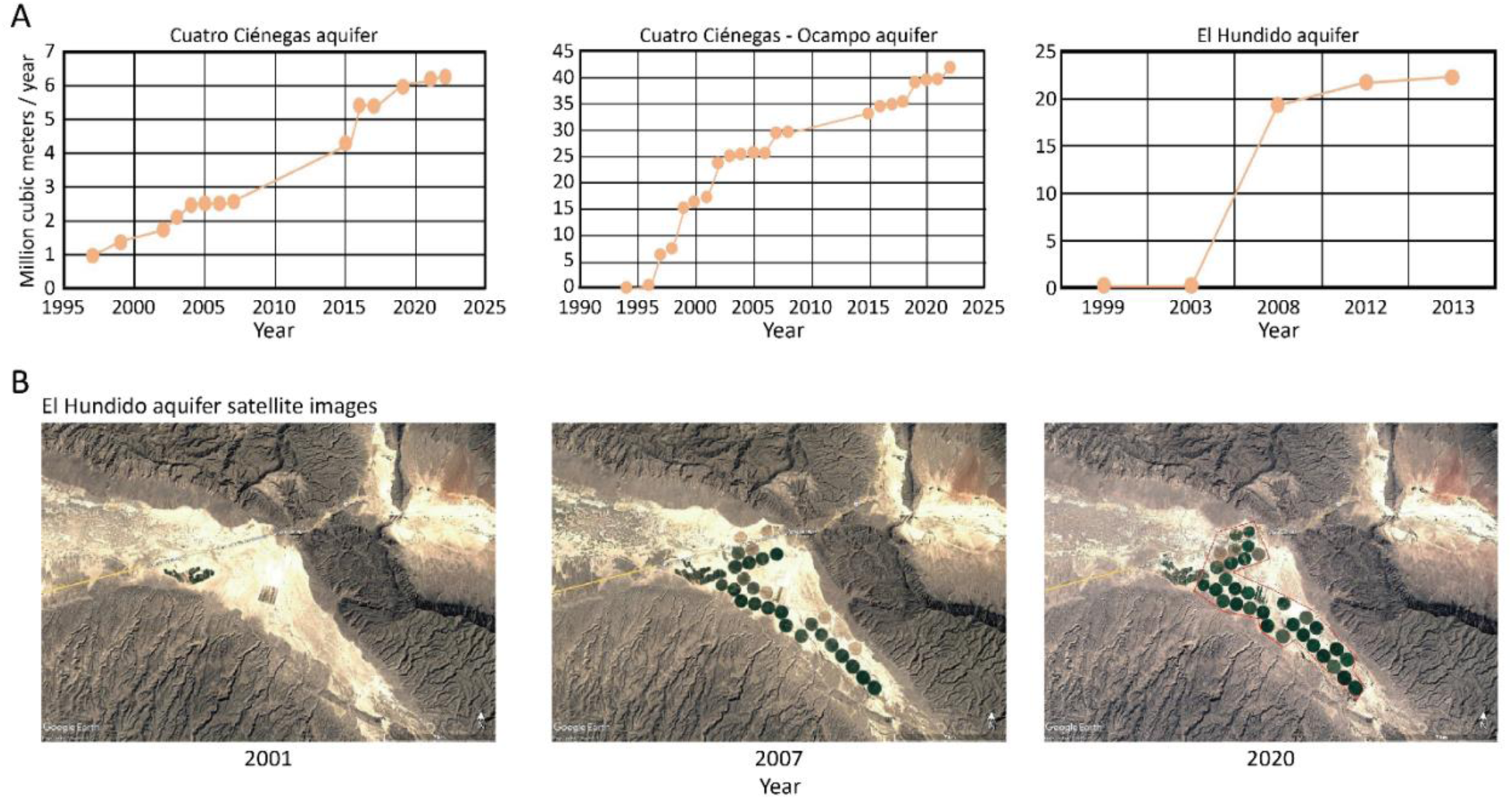
Expansion of alfalfa cultivation and increasing water extraction from Cuatro Ciénegas aquifers. (A) Annual water consumption for agricultural practices (millions of cubic meters per year) in three aquifers: Cuatro Ciénegas, Ocampo, and El Hundido (data from IMTA). (B) Google Earth images from 2001, 2007, and 2020 showing circular alfalfa fields (rondines) in the Hundido Valley. At these time points, 21 rondines were active in 2007 and 29 in 2020, illustrating the expansion of alfalfa cultivation.

**Supplementary Fig. 2.**
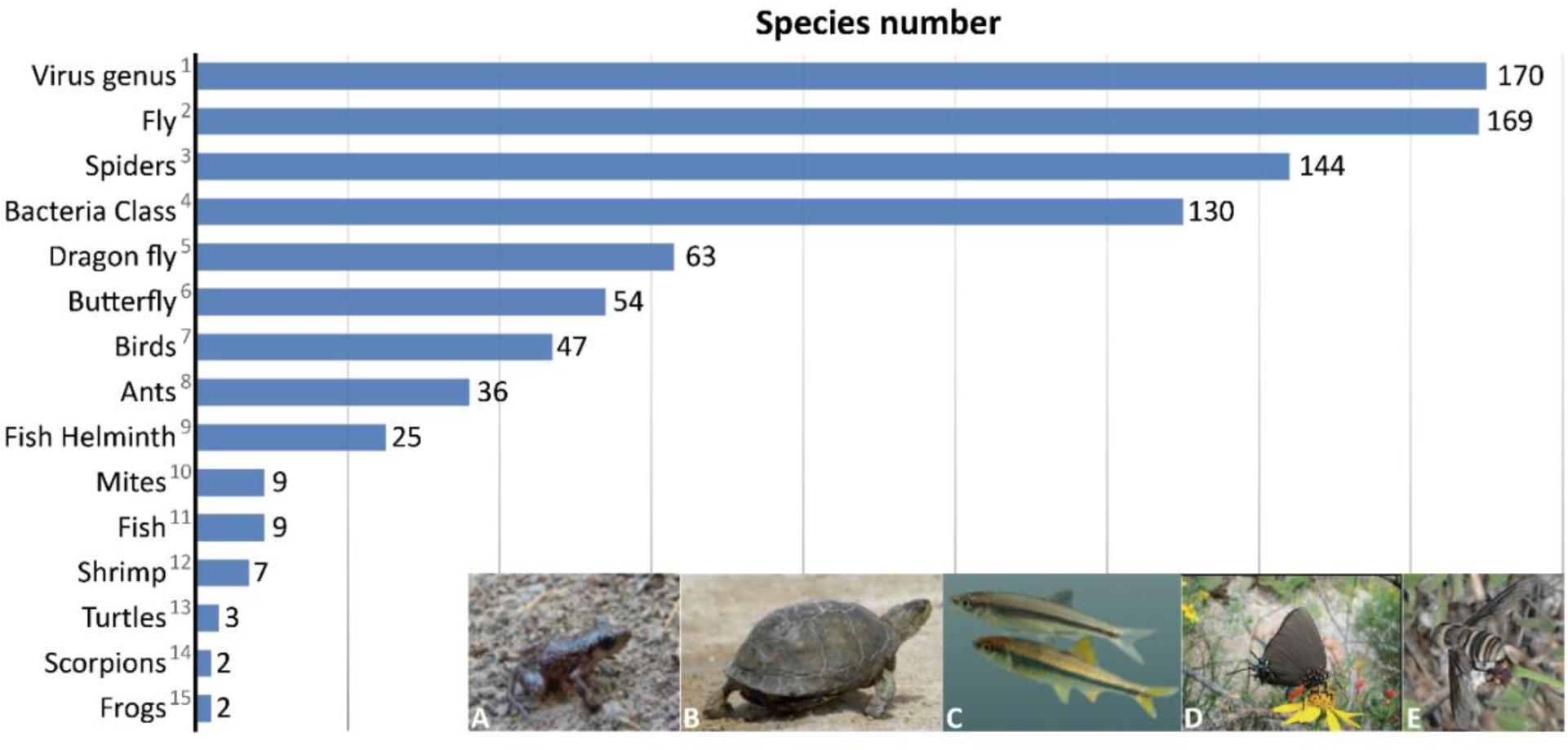
Inventory of endemic biodiversity in the Cuatro Ciénegas Basin (2013). The chart summarizes the number of species comprising the basin’s unique biodiversity. Representative photographs include: (A) *Eleutherodactylus* sp. nov. (frog), (B) T*errapene coahuila* (turtle), (C) *Cyprinella xanthicara* (fish), (D) *Atlides halesus* (butterfly; Cramer), and (E) *Exoprosopa* sp. (bee). A reference is provided to the publication where detailed results on the diversity of viruses, bacteria, mites, fish, frogs, and other organisms were previously reported. References: 1.- Isa, P. *et al*., 2018, 2.- Ávalos-Hernández *et al*., 2019, 3.- Bizuet-Flores *et al*., 2015, 4.- Souza *et al*., 2018, 5.- Ortega-Salas, H., and González-Soriano, E., 2019, 6.- Hernández-Jerónimo *et al*., 2019, 7.- Corcuera *et al*., 2019, 8.- Janda *et al*., 2019, 9.- Pérez-Ponce de León, G., and Aguilar-Aguilar, R., 2019, 10.- Paredes-León, R., 2019, 11.- Espinosa-Pérez, H., and Lambarri-Martínez, C. 2019, 12.- García-Vázquez *et al*., 2022, 13.- García-Vázquez *et al*., 2021, 14.- Francke B., O.F. 2019, 15.- García-Vázquez *et al*., 2019.

**Supplementary Fig. 3.**
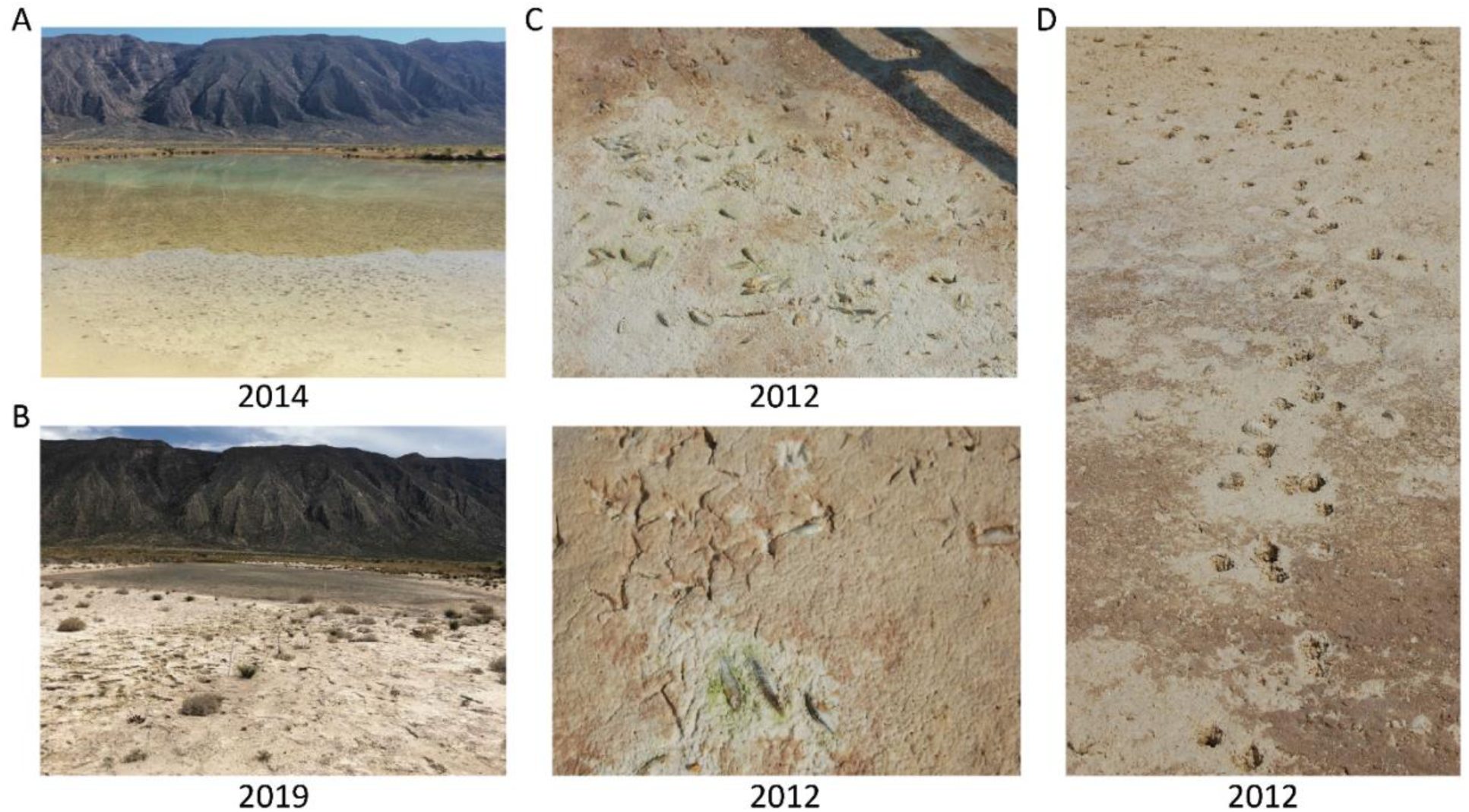
Ecological impacts of desiccation in the Churince Intermediate Lagoon. Photographs show: (A) surface water still present in 2014; (B) by 2019, water restricted to the lagoon’s center; (C) dead fish and bird footprints on the exposed lagoon bed; and (D) turtle tracks indicating individuals followed the receding water and likely perished.

**Supplementary Fig. 4.**
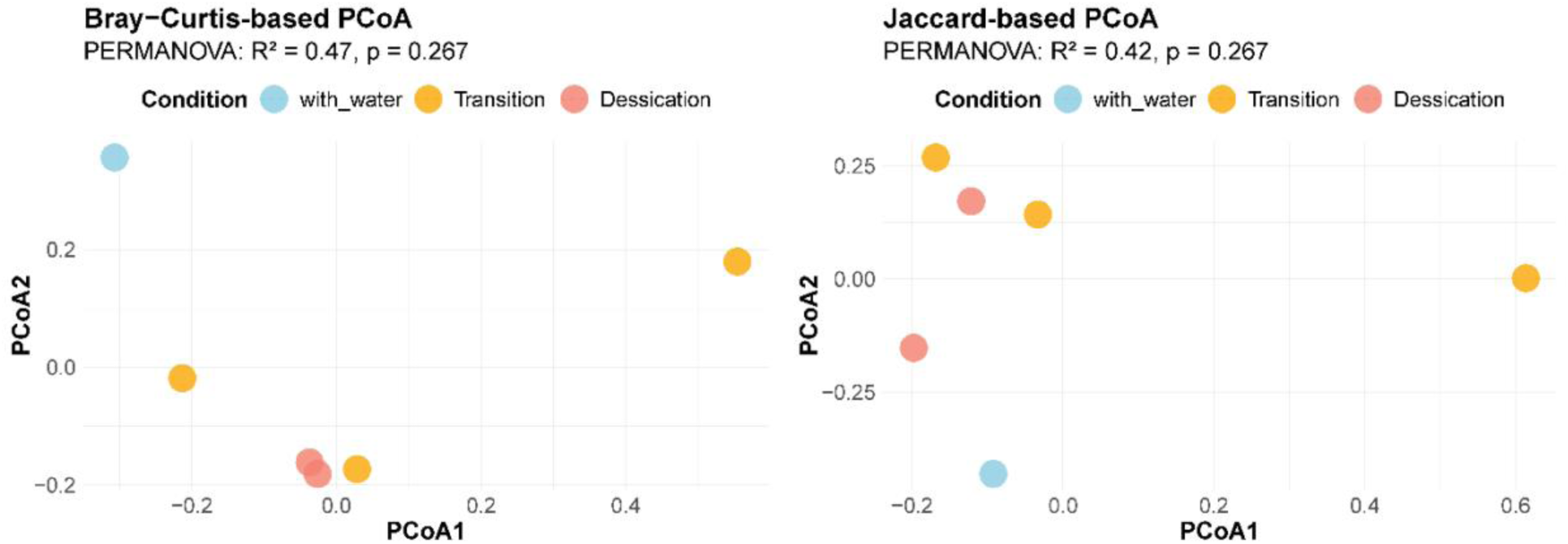
Beta diversity of culturable bacterial communities across ecological phases. (A) Principal Coordinates Analysis (PCoA) based on Bray–Curtis dissimilarity, illustrating clustering of bacterial communities by ecological phase (With Water, Transition, and Desiccation). PERMANOVA results (R² and p-values) are shown. (B) PCoA based on Jaccard dissimilarity, showing community clustering by hydrological condition. PERMANOVA results (R² and p-values) are indicated.

**Supplementary Fig. 5.**
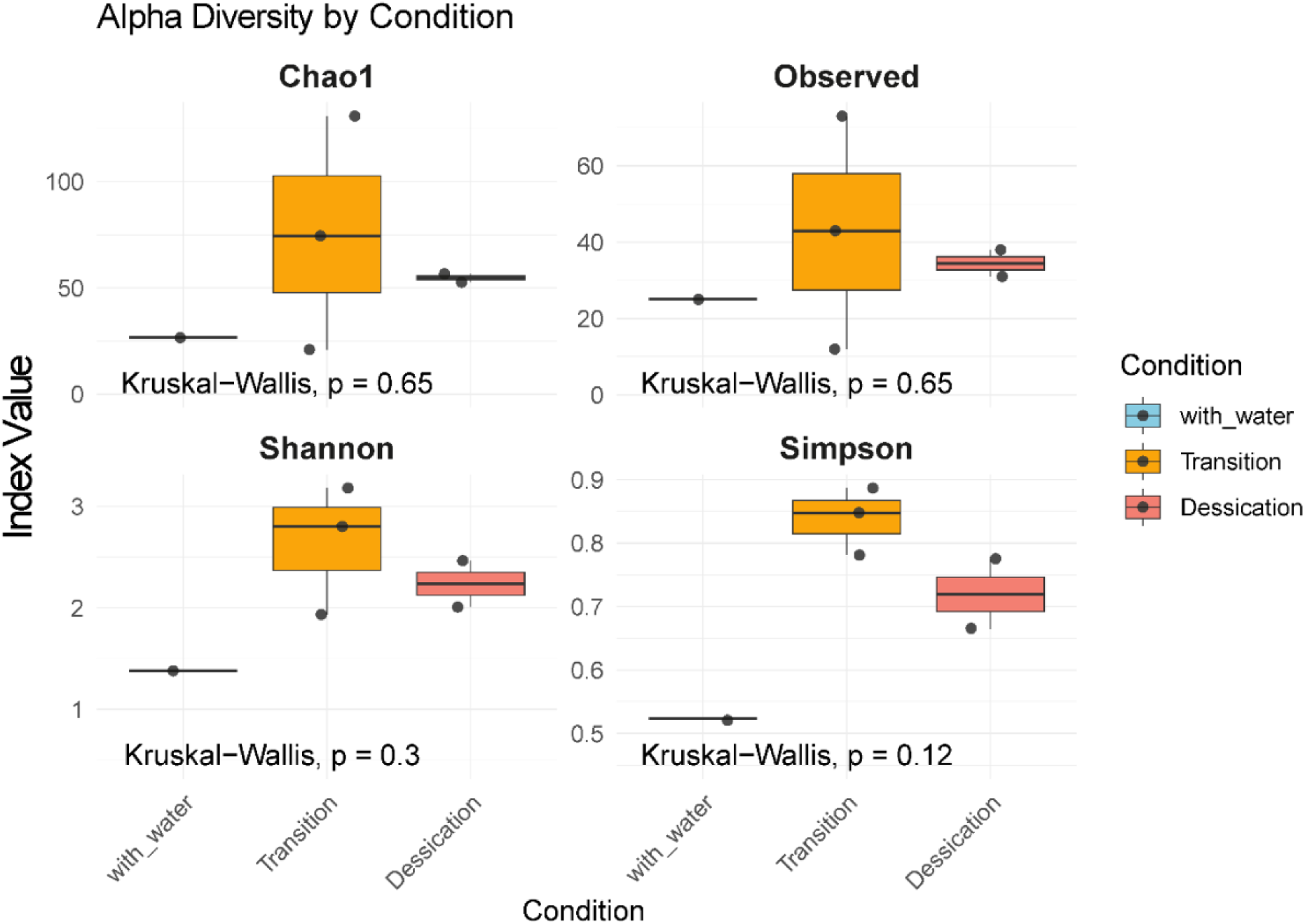
Alpha diversity of culturable bacterial communities across ecological phases. Box plots show the distribution of alpha diversity indices for bacterial isolates from Churince Intermediate Lagoon sediments across three ecological phases: With Water, Transition, and Desiccation. (A) Chao1 richness estimator. (B) Observed species index. (C) Shannon diversity index. (D) Simpson diversity index. Statistical comparisons among phases were performed using the Kruskal–Wallis test; p-values are shown on each plot.

**Supplementary Fig. 6.**
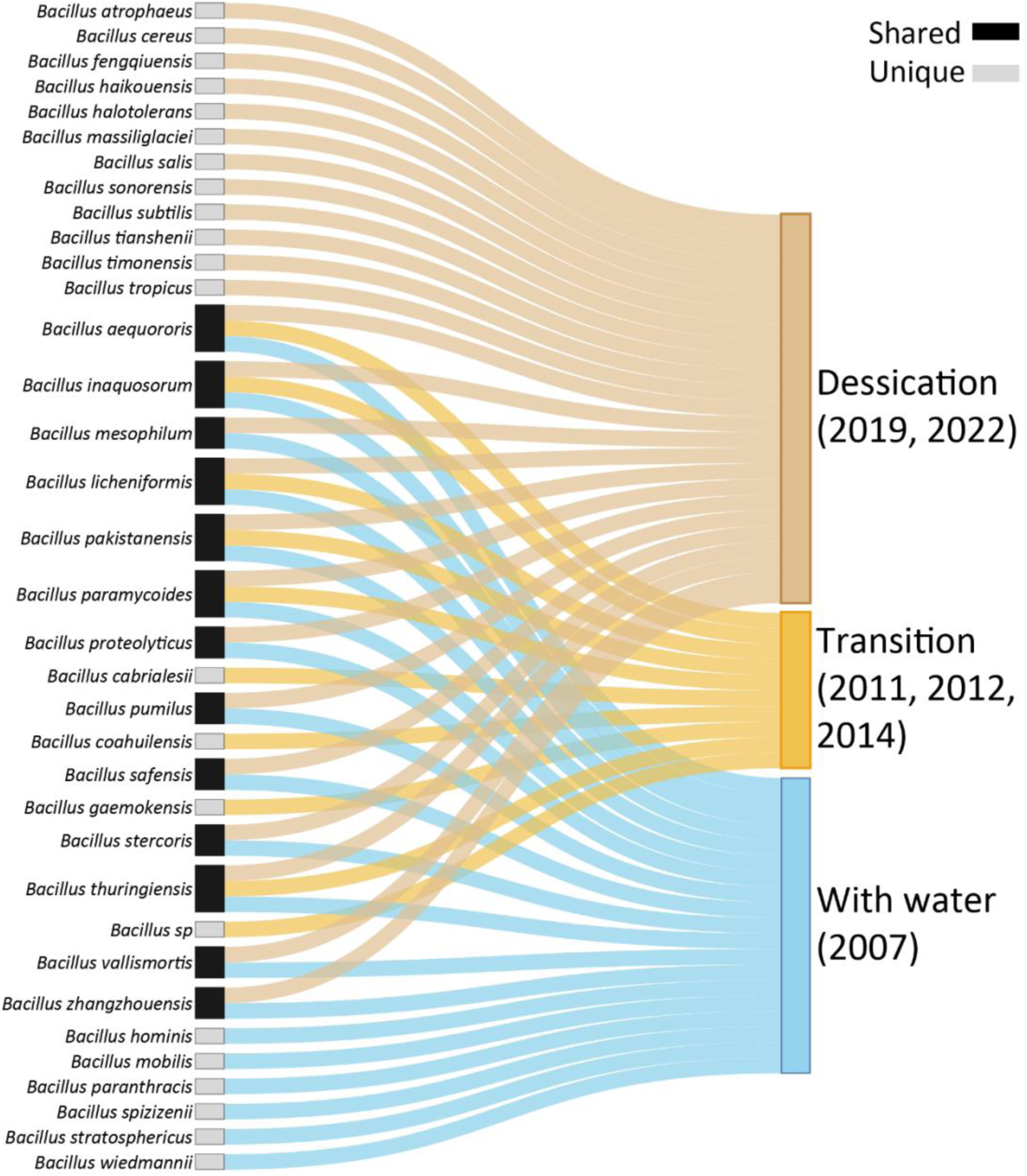
Distribution of culturable *Bacillus* species across ecological phases in sediments of the Churince Intermediate Lagoon. The Sankey diagram illustrates the presence and overlaps of *Bacillus* species identified from culturable isolates collected during three ecological phases: With Water (2007 sampling), Transition (intermediate desiccation; 2011, 2012, and 2014 sampling), and Desiccation (fully dry; 2019 and 2022 sampling). Grey boxes denote species unique to a single phase, whereas black boxes represent species occurring in at least two phases.

**Supplementary Fig. 7.**
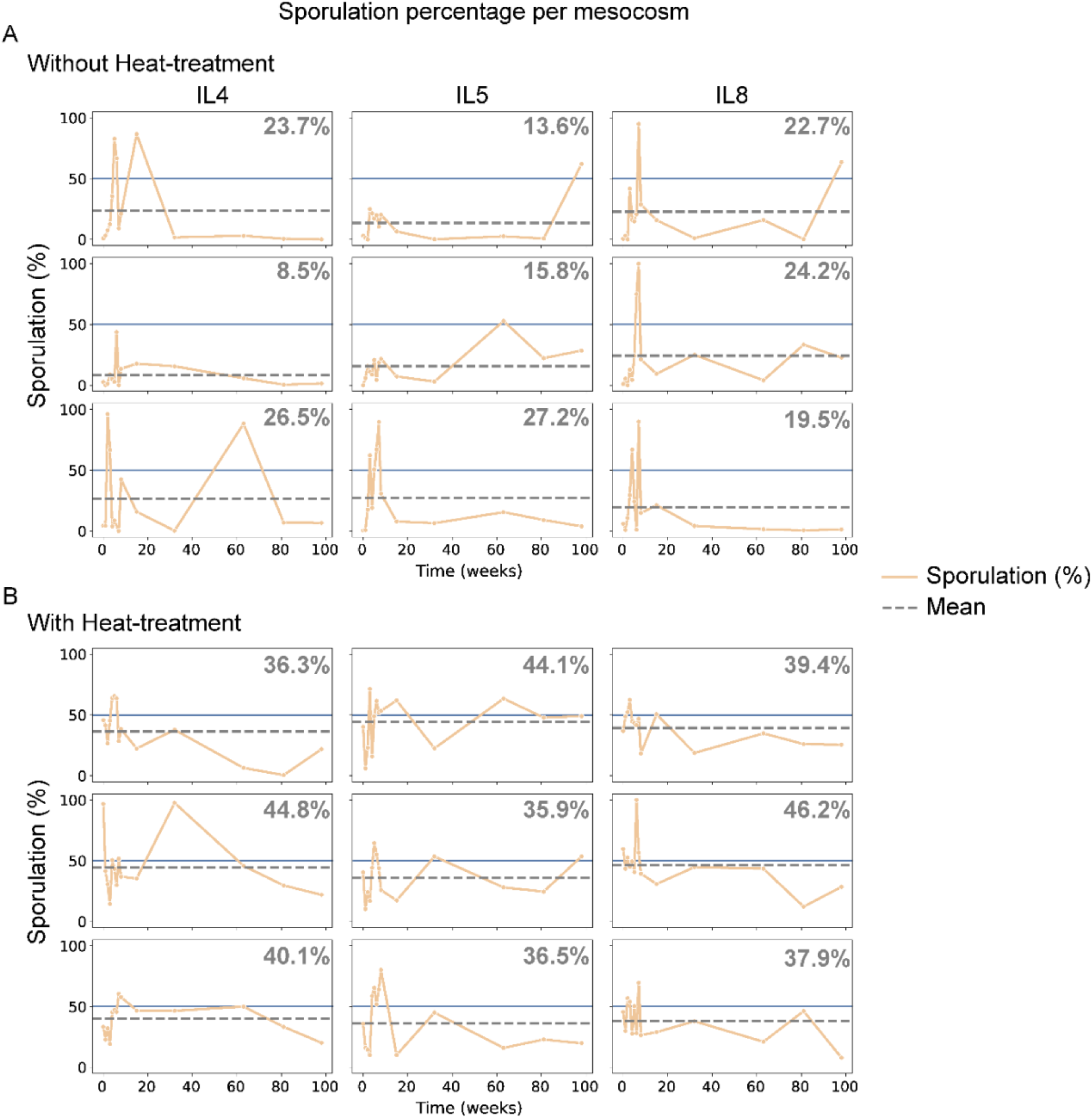
Sporulation dynamics across mesocosms. Line plots show temporal variation in sporulation percentage (%) over time (weeks) for individual mesocosms. Each panel corresponds to a single mesocosm. Orange lines indicate observed values, and dashed gray lines show the mean sporulation level across all time points, with mean sporulation (%) across all sampling times shown in the top right of each plot. Columns represent samples from sites IL4, IL5, and IL8 (see Fig. 6), each with three independent replicates. Panel (A) shows mesocosms initiated without prior heat treatment; panel (B) shows mesocosms initiated with heat treatment.

## Supplementary Tables

**Supplementary Table 1.** Coordinates, CFU counts, and sporulation percentages from Laguna Intermedia (Churince) sampling sites.

**Supplementary Table 2.-** CFU counts from transect sampling in the Churince system (2011).

**Supplementary Table 3.-** Morphological characterization of colony phenotypes from a 50- cm sediment core at the LI7 sampling site.

**Supplementary Table 4.-** Colony Count and Percentage of Spores from vertical-sectional study at the LI7 Sampling Site.

**Supplementary Table 5.-** Monthly groundwater levels of the Intermediate Lagoon (2007– 2015).

**Supplementary Table 6.-** Taxonomic identification of 1,419 culturable bacterial isolates from the Churince hydrological system (2007–2022).

**Supplementary Table 7.**- Relative and absolute abundances (n) of culturable bacterial genera isolated from the Churince hydrological system.

**Supplementary Table 8.-** Relative and absolute abundances (n) of ***Bacillus*** spp. across ecological phases defined by water availability.

**Supplementary Table 9.-** Kruskal–Wallis test of *Bacillus* spp. across ecological phases defined by water availability.

**Supplementary Table 10.-** Colony count and spore percentage from mesocosms

## Bibliography

[1] . Abdallah, M. Ben, Karray, F., Mhiri, N., Mei, N., Quéméneur, M., Cayol, J.-L., Erauso, G., Tholozan, J.-L., Alazard, D., & Sayadi, S. (2016). Prokaryotic diversity in a Tunisian hypersaline lake, Chott El Jerid. Extremophiles, 20(2), 125–138. 10.1007/s00792-015-0805

[2] . Alcaraz, L. D., Olmedo, G., Bonilla, G., Cerritos, R., Hernández, G., Cruz, A., Ramírez, E., Putonti, C., Jiménez, B., Martínez, E., López, V., Arvizu, J. L., Ayala, F., Razo, F., Caballero, J., Siefert, J., Eguiarte, L., Vielle, J. P., Martínez, O., Souza, V., Herrera-Estrella, A., & Herrera-Estrella, L. (2008). The genome of Bacillus coahuilensis reveals adaptations essential for survival in the relic of an ancient marine environment. Proceedings of the National Academy of Sciences of the United States of America, 105(15), 5803–5808. 10.1073/pnas.0800981105

[3] . Allison, S. D., & Martiny, J. B. H. (2008). Resistance, resilience, and redundancy in microbial communities. Proceedings of the National Academy of Sciences, 105(supplement_1), 11512–11519. 10.1073/pnas.0801925105

[4] . Bierkens, M. F. P., & Wada, Y. (2019). Non-renewable groundwater use and groundwater depletion: a review. Environmental Research Letters, 14(6), 063002. 10.1088/1748-9326/ab1a5f

[5] . Brown, P. D., Schröder, T., Ríos-Arana, J. V., Rico-Martinez, R., Silva-Briano, M., Wallace, R. L., & Walsh, E. J. (2022). Processes contributing to rotifer community assembly in shallow temporary aridland waters. Hydrobiologia, 849(17–18), 3719–3735. 10.1007/s10750-022-04842-8

[6] . Cerritos, R., Eguiarte, L. E., Avitia, M., Siefert, J., Travisano, M., Rodríguez-Verdugo, A., & Souza, V. (2011). Diversity of culturable thermo-resistant aquatic bacteria along an environmental gradient in Cuatro Ciénegas, Coahuila, México. Antonie van Leeuwenhoek, International Journal of General and Molecular Microbiology, 99(2), 303–318. 10.1007/s10482-010-9490-9

[7] . Chanal, A., Chapon, V., Benzerara, K., Barakat, M., Christen, R., Achouak, W., Barras, F., & Heulin, T. (2006). The desert of Tataouine: an extreme environment that hosts a wide diversity of microorganisms and radiotolerant bacteria. Environmental Microbiology, 8(3), 514–525. 10.1111/j.1462-2920.2005.00921.x

[8] . Davidson, N. C., Finlayson, C. M., McInnes, R. J., Rostron, C., Simpson, M., & Gell, P. A. (2023). What’s happening to the world’s wetlands? In Ramsar Wetlands (pp. 219–235). Elsevier. 10.1016/B978-0-12-817803-4.00019-X

[9] . Filippidou, S., Wunderlin, T., Junier, T., Jeanneret, N., Dorador, C., Molina, V., Johnson, D. R., & Junier, P. (2016). A Combination of Extreme Environmental Conditions Favor the Prevalence of Endospore-Forming Firmicutes. Frontiers in Microbiology, 7. 10.3389/fmicb.2016.01707

[10] . Gasbarro, R., Chu, J. W. F., & Tunnicliffe, V. (2019). Disassembly of an epibenthic assemblage in a sustained severely hypoxic event in a northeast Pacific basin. Journal of Marine Systems, 198, 103184. 10.1016/j.jmarsys.2019.103184

[11] . Gilbert, J. A., Peixoto, R. S., Scholz, A. H., Dominguez Bello, M. G., Korsten, L., Berg, G., Singh, B., Boetius, A., Wang, F., Greening, C., Wrighton, K., Bordenstein, S., Jansson, J. K., Lennon, J. T., Souza, V., Thomas, T., Cowan, D., Crowther, T. W., Nguyen, N., Harper, L., Haraoui, L.-P., Ishaq, S. L., & Redford, K. (2025). Launching the IUCN Microbial Conservation Specialist Group as a global safeguard for microbial biodiversity. Nature Microbiology. 10.1038/s41564-025-02113-5

[12] . Haglund, A.-L., Lantz, P., TÃ¶rnblom, E., & Tranvik, L. (2003). Depth distribution of active bacteria and bacterial activity in lake sediment. FEMS Microbiology Ecology, 46(1), 31–38. 10.1016/S0168-6496(03)00190-9

[13] . Higgins, D., & Dworkin, J. (2012). Recent progress in *Bacillus subtilis* sporulation. FEMS Microbiology Reviews, 36(1), 131–148. 10.1111/j.1574-6976.2011.00310.x

[14] . Kheiri, R., Mehrshad, M., Pourbabaee, A. A., Ventosa, A., & Amoozegar, M. A. (2023). Hypersaline Lake Urmia: a potential hotspot for microbial genomic variation. Scientific Reports, 13(1), 374. 10.1038/s41598-023-27429-2

[15] . Lennon, J. T., & Jones, S. E. (2011). Microbial seed banks: the ecological and evolutionary implications of dormancy. Nature Reviews Microbiology, 9(2), 119–130. 10.1038/nrmicro2504

[16] . Letunic, I., & Bork, P. (2024). Interactive Tree of Life (iTOL) v6: recent updates to the phylogenetic tree display and annotation tool. Nucleic Acids Research, 52(W1), W78–W82. 10.1093/nar/gkae268

[17] . Leung, P. M., Bay, S. K., Meier, D. V., Chiri, E., Cowan, D. A., Gillor, O., Woebken, D., & Greening, C. (2020). Energetic Basis of Microbial Growth and Persistence in Desert Ecosystems. MSystems, 5(2). 10.1128/mSystems.00495-19

[18] . McDonald, M. D., Owusu-Ansah, C., Ellenbogen, J. B., Malone, Z. D., Ricketts, M. P., Frolking, S. E., Ernakovich, J. G., Ibba, M., Bagby, S. C., & Weissman, J. L. (2024). What is microbial dormancy? Trends in Microbiology, 32(2), 142–150. 10.1016/j.tim.2023.08.006

[19] . O’Neill, B. J. (2016). Community disassembly in ephemeral ecosystems. Ecology, 97(12), 3285–3292. 10.1002/ecy.1604

[20] . Ortiz Acosta, S. E., & Romo Aguilar, M. de L. (2016). Impactos socioambientales de la gestión del agua en el área natural protegida de Cuatro Ciénegas, Coahuila. Región Y Sociedad, 28(66). 10.22198/rys.2016.66.a405

[21] . Ostfeld, R. S., & LoGiudice, K. (2003). Community Disassembly, Biodiversity Loss, and the Erosion of an Ecosystem Service. Ecology, 84(6), 1421–1427. http://www.jstor.org/stable/3107961

[22] . Pisanty, I., Rodríguez-Sánchez, M., Peralta-García, C., Cervantes-Campero, G., Souza, V., & Mandujano, M. C. (2022). Plants as a Canary in the Mine: A Wetland Response to Ecosystem Failure (pp. 121–142). 10.1007/978-3-030-83270-4_8

[23] . Piubeli, F., de Lourdes Moreno, M., Kishi, L. T., Henrique-Silva, F., García, M. T., & Mellado, E. (2015). Phylogenetic Profiling and Diversity of Bacterial Communities in the Death Valley, an Extreme Habitat in the Atacama Desert. Indian Journal of Microbiology, 55(4), 392–399. 10.1007/s12088-015-0539-3

[24] . Prosser, J. I., Bohannan, B. J. M., Curtis, T. P., Ellis, R. J., Firestone, M. K., Freckleton, R. P., Green, J. L., Green, L. E., Killham, K., Lennon, J. J., Osborn, A. M., Solan, M., van der Gast, C. J., & Young, J. P. W. (2007). The role of ecological theory in microbial ecology. Nature Reviews Microbiology, 5(5), 384–392. 10.1038/nrmicro1643

[25] . Ramos-Madrigal, C., Martínez-Romero, E., Tapia-Torres, Y., & Servín-Garcidueñas, L. E. (2024). Bacterial Diversity Profiles of Desert Sand and Salt Crusts from the Gran Desierto de Altar, Sonora, Mexico. Diversity, 16(12), 745. 10.3390/d16120745

[26] . Rodrigues, C. J. C., & de Carvalho, C. C. C. R. (2022). Cultivating marine bacteria under laboratory conditions: Overcoming the “unculturable” dogma. Frontiers in Bioengineering and Biotechnology, 10. 10.3389/fbioe.2022.964589

[27] . Rodríguez-Torres, D. M., Islas-Robles, Á., Gómez-Lunar, Z., Delaye, L., Hernández-González, I., Souza, V., Travisano, M., & Olmedo-Álvarez, G. (2017). Phenotypic Microdiversity and Phylogenetic Signal Analysis of Traits Related to Social Interaction in Bacillus spp. from Sediment Communities. Frontiers in Microbiology, 8. 10.3389/fmicb.2017.00029

[28] . Scheffer, M., Carpenter, S., Foley, J. A., Folke, C., & Walker, B. (2001). Catastrophic shifts in ecosystems. Nature, 413(6856), 591–596. 10.1038/35098000

[29] . Scheffer, M., Carpenter, S. R., Lenton, T. M., Bascompte, J., Brock, W., Dakos, V., van de Koppel, J., van de Leemput, I. A., Levin, S. A., van Nes, E. H., Pascual, M., & Vandermeer, J. (2012). Anticipating Critical Transitions. Science, 338(6105), 344–348. 10.1126/science.1225244

[30] . Schimel, J. P. (2018). Life in Dry Soils: Effects of Drought on Soil Microbial Communities and Processes. *Annual Review of Ecology*, Evolution, and Systematics, 49(1), 409–432. 10.1146/annurev-ecolsys-110617-062614

[31] . Setlow, P. (2014). Germination of Spores of Bacillus Species: What We Know and Do Not Know. Journal of Bacteriology, 196(7), 1297–1305. 10.1128/JB.01455-13

[32] . Setlow, P., & Christie, G. (2023). New Thoughts on an Old Topic: Secrets of Bacterial Spore Resistance Slowly Being Revealed. Microbiology and Molecular Biology Reviews, 87(2). 10.1128/mmbr.00080-22

[33] . Shoemaker, W. R., & Lennon, J. T. (2018). Evolution with a seed bank: The population genetic consequences of microbial dormancy. Evolutionary applications, 11(1), 60–75. 10.1111/eva.12557

[34] . Souza, V., Espinosa-Asuar, L., Escalante, A. E., Eguiarte, L. E., Farmer, J., Forney, L., Lloret, L., Rodríguez-Martínez, J. M., Soberón, X., Dirzo, R., & Elser, J. J. (2006). An endangered oasis of aquatic microbial biodiversity in the Chihuahuan desert. Proceedings of the National Academy of Sciences, 103(17), 6565–6570. 10.1073/pnas.0601434103

[35] . Souza, V., Moreno-Letelier, A., Travisano, M., Alcaraz, L. D., Olmedo, G., & Eguiarte, L. E. (2018). The lost world of cuatro ciénegas basin, a relictual bacterial niche in a desert oasis. ELife, 7, 1–17. 10.7554/eLife.38278

[36] . Stulina, G., Verkhovtseva, N., & Gorbacheva, M. (2019). Composition of the Microorganism Community Found in the Soil Cover on the Dried Seabed of the Aral Sea. Journal of Geoscience and Environment Protection, 07(08), 1–23. 10.4236/gep.2019.78001

[37] . Vreeland, R. H., Rosenzweig, W. D., & Powers, D. W. (2000). Isolation of a 250 million-year-old halotolerant bacterium from a primary salt crystal. Nature, 407(6806), 897–900. 10.1038/35038060

[38] . Wang, G., & Or, D. (2013). Hydration dynamics promote bacterial coexistence on rough surfaces. The ISME Journal, 7(2), 395–404. 10.1038/ismej.2012.115

[39] . Zavaleta, E., Pasari, J., Moore, J., Hernández, D., Suttle, K. B., & Wilmers, C. C. (2009). Ecosystem Responses to Community Disassembly. Annals of the New York Academy of Sciences, 1162(1), 311–333. 10.1111/j.1749-6632.2009.04448.x

